# Multi-omic dissection of ancestral heat stress memory responses in *Brachypodium distachyon*

**DOI:** 10.1101/2023.03.04.531132

**Authors:** Zheng Xinghai, Qiao Wen Tan, Peng Ken Lim, Marek Mutwil

## Abstract

Stressful environmental conditions, including heat stress (HS), are a major limiting factor in crop yield. Understanding the molecular mechanisms of plant stress memory and resilience is important for engineering more resistant plants and improving crop yield. To study how the different gene regulatory layers change upon repeated HS and how these layers are interconnected, we performed a dense temporal atlas of gene expression, alternative splicing, small and long noncoding RNAs, and DNA methylation in *Brachypodium distachyon*. Results show that a second HS induces changes in coding and noncoding RNA expression and alternative splicing and that DNA demethylation is responsible for mediating differential gene expression. We identified a long noncoding RNA regulatory network and provided evidence that lncRNAs positively regulate gene expression, while miRNAs are implicated in alternative splicing events. We reconstructed the ancestral heat memory network of flowering plants by comparing the dynamic responses of *Arabidopsis thaliana* and *Brachypodium distachyon*. These findings enhance our understanding of the complex inter-layer cross-talk governing HS resilience and memory and identify novel genes essential for these processes.

## Introduction

Organisms are regularly subjected to stressful environmental conditions, which impede the organism’s development and growth and are known as stress. Climate change is expected to result in more extreme weather and more frequent heat stress (HS)(Battisti and Naylor, 2009; Lobell et al., 2011), and HS is one of the major limiting factors in crop yield. Consequently, elucidating the mechanisms by which plants respond to high temperature and acclimatize to future stresses are of great interest. All organisms exhibit innate thermotolerance to heat stress (basal tolerance). Still, they also can acquire tolerance to otherwise lethal HS (acquired tolerance)(Larkindale et al., 2005), by being exposed to non-lethal HS. The acquired thermotolerance can persist for several days (Richter et al., 2010; Asensi-Fabado et al., 2017; Ohama et al., 2017; Li et al., 2017b), which requires lasting modifications to gene activity and metabolism (Asensi-Fabado et al., 2017; Olas et al., 2021). This somatic stress’ memory’ has been studied for stresses such as heat (Charng et al., 2006; Charng et al., 2007; Stief et al., 2014; Brzezinka et al., 2016; Brzezinka et al., 2019; Friedrich et al., 2021), drought (Ding et al., 2012; Kim et al., 2012), salt (Feng et al., 2016), and immunity (Jaskiewicz et al., 2011; Singh et al., 2014; Mozgová et al., 2015).

HS memory has been observed on multiple layers of gene expression regulation. HS induces the expression of heat shock transcription factors (HSF) and heat shock proteins (HSP) that show sustained expression for days after the stress has passed (Sedaghatmehr et al., 2022). HS has been shown to repress alternative splicing, but to a lesser degree for plants that were exposed to a prior HS (Ling et al., 2018). Noncoding RNAs, including microRNAs (miRNAs)(Giacomelli et al., 2012; Guan et al., 2013; Stief et al., 2014; Zhao et al., 2016), small interfering RNAs (siRNAs)(Ito et al., 2011; Yu et al., 2013; Li et al., 2014), and long noncoding RNAs (lncRNAs)(Wang et al., 2019) are known to play a role in heat stress response and memory (Zhao et al., 2020). Plants also undergo extensive epigenetic reprogramming to resist HS. DNA methylation is thought to regulate genes essential for HS response (Popova et al., 2013; Liu et al., 2015; Liu et al., 2017), as the global DNA methylation increased in *Arabidopsis thaliana* and *Quercus suber* during HS (Boyko et al., 2010; Correia et al., 2013), but decreased in strawberry and in another Arabidopsis study (Korotko et al., 2021; López et al., 2022). RNA-directed DNA methylation (RdDM) is required for basal thermotolerance, and plants deficient in RdDM are hypersensitive to HS (Zhao et al., 2020). DNA methylation affects the expression of HS-induced genes, and plants with deficient CHH methylation have an improved HS tolerance (Shen et al., 2014b). Histone modifications are also crucial for thermotolerance (Zhao et al., 2020), as plants deficient in *histone acetyltransferase GCN5* or *histone deacetylase 6* (*HDA6*) exhibit severe defects in thermotolerance under HS (Hu et al., 2015). Finally, HS has been shown to cause chromatin remodeling and change the 3D structure of the genome by altering the large-scale chromatin interactions (Yadav et al., 2022).

HS resilience and memory thus require a complex, dynamic interplay between the different gene regulatory layers (Zhao et al., 2020), but our knowledge of how these layers interact and control gene expression in more than a few species is limited. To address this, we performed a dense temporal atlas capturing gene expression, alternative splicing, small and long noncoding RNAs, and DNA methylation. In addition, a comprehensive statistical analysis revealed the genes and biological pathways affected by the changes in these regulatory layers and identified potential functional inter-layer associations.

Here, we show that a second HS induces large changes to coding and noncoding RNA expression and alternative splicing in the model grass *Brachypodium distachyon*. We observed a significant DNA demethylation event after the first HS and showed that this is connected to mediating differential gene expression in the second HS. We identified a long noncoding RNA and lncRNA-mRNA interaction network and provided evidence that lncRNAs positively regulate gene expression. The analysis of miRNAs revealed their role in alternative splicing events. Finally, we reconstructed the ancestral heat memory network of flowering plants by comparing the dynamic responses of *Arabidopsis* and *Brachypodium*. These findings not only shed light on the complex inter-layer cross-talk governing HS resilience and memory but also provide information on novel genes important for these processes. We envision our findings in the model grass *Brachypodium* will help engineer more heat-tolerant crops.

## Methods

### Brachypodium distachyon growth conditions

Four *B. distachyon* Bd21-3 seeds were sowed in the soil and vernalized for 5 days in the 4 cold room. After vernalization, the seeds were grown at 25, 50% relative humidity, 100µE*m^-2^*s^-1^ light intensity, and with a day/night cycle of 20/4 hours.

### Acclimation to lethal heat stress experiment

Twenty-one days after vernalization, *B. distachyon* Bd21-3 seedlings were divided into acclimated and non-acclimated groups. Before a 90-minute heat stress treatment, the acclimated group underwent a 1-day 39L heat priming and a 2-day recovery, while the non-acclimated group was kept at 25L for three days. In the 90-minute heat stress treatment, the acclimated and non-acclimated plants were divided into 42L, 48L, and, 54L sub-groups. After the heat stress treatment, the plants recovered at 25L temperature for four days. The images of three biological replicates at five time points were recorded.

### Acclimation experiment and sample collection

*B. distachyon* Bd21-3 seedlings were divided into an acclimated group and a non-acclimated group after 21 days from vernalization. Before a 1-day 39 heat stress treatment, the acclimated group underwent a 1-day 39L heat priming and a 2-day recovery, while the non-acclimated group was kept at 25 for three days. In each group, leaves of 3 biological replicates at the ten time points were cut and immediately stored in a −80L freezer.

### RNA extraction and sequencing

Total RNA was extracted from samples using Spectrum™ Plant Total RNA Kit according to the manufacturer’s protocol. Novogene conducted the preparation of RNA library and transcriptome sequencing. After receiving raw data from Novogene, adapters and low-quality reads were filtered by Trimmomatic v0.38 (Bolger et al., 2014). Trimmed reads were then aligned two times against the reference genome (*Brachypodium distachyon* Bd21-3 v1.2) from Phytozome v13 (http://www.phytozome.net) (Goodstein et al., 2012) using TopHat v2.1.1 (Kim et al., 2013). The second alignment was done to improve the reads mapping around spliced sites using the information of junction sites generated from the first alignment.

### mRNA analysis

With the mapping file (.bam) from the second alignment as a base, reads were counted by featureCounts v2.0.1 (Liao et al., 2014) mapping the (.bam) file to the (.gtf) file downloaded from Phytozome. After constructing the count matrix, genes with zero counts across all samples were removed. Subsequently, differentially expressed analysis was performed in DESeq2 v1.26.0 (Love et al., 2014), with Benjamini-Hochberg correction (Benjamini and Hochberg, 1995). Differentially expressed mRNAs (DEmRNAs) were selected based on the absolute value of log2FoldChange >= 1 and the adjusted p-value < 0.05. Next, TPM values were calculated based on the gene count values and their respective lengths. After constructing the TPM matrix, PCA and k-means cluster analyses were performed.

### Alternative splicing analysis

Alternative splicing (AS) events were counted by rMATs v4.1.1 (Shen et al., 2014a) using the mapping (.bam) file and (.gtf) file downloaded from Phytozome. Alternative splicing genes (ASGs) and differential alternative splicing transcripts (DASTs) were selected from the rMATs result file (*.MATS.JC.txt) for cases where the FDR value was less than 0.05.

Given that the size and composition of ASGs remain consistent across comparisons, while the types of ASGs vary, we grouped the ASGs in all AllvsAA1 comparisons together as a background, separated by different types (Figure 3A & S7). The relationship between mRNA expression and alternative splicing was determined by performing a hypergeometric test on comparisons between differentially expressed genes (DEGs) and differentially alternatively spliced transcripts (DASTs).

### Long noncoding RNA analysis

Mapping files (.bam) were processed using StringTie v2.1.7 (Pertea et al., 2015), which assembles the whole transcripts set for each sample. The assembled transcriptomes were then merged in a unique reference transcriptome with Cuffmerge, and transcript sequences were extracted with Gffread v0.12.7 (Pertea and Pertea, 2020). Transcript sequences were subjected to five consecutive filters to identify lncRNAs (De Quattro et al., 2018): (i) length ≥ 200 bp; (ii) ORF ≤100 amino acids; (iii) no homology with known protein domain; (iv) a low coding potential; (v) no homology with structural RNAs. Next, we generated a .gtf file for the found lncRNAs and used the mapping file (.bam) from the second alignment to count lncRNA reads by featureCounts v2.0.1. After the count matrix was built, differential lncRNA expression analysis was performed as for mRNA.

### miRNA extraction and miRNA-Seq analysis

miRNA was extracted from samples using mirVana™ PARIS™ RNA and Native Protein Purification Kit according to the manufacturer’s protocol and were sent to Novogene for miRNA Sequencing. Novogene conducted the preparation of the miRNA library and microRNA sequencing. After receiving raw data from Novogene, the alignment of the cleaned reads to mature miRNAs downloaded from mirBase (Kozomara et al., 2019) was done using the short read aligner Bowtie v1.0.0 (Langmead, 2010). Reads with no more than two mismatches in the alignment to the mature miRNAs were selected. Counts were extracted by Samtools v1.15.1 (Li et al., 2009), and the differential expression analysis was done as for mRNA and lncRNA.

### DNA extraction and DNA-methylation analysis

DNA was extracted from samples using the CTAB method and was sent to Novogene for Whole Genome Bisulfite Sequencing (WGBS). Novogene conducted the preparation of the DNA library and Whole Genome Bisulfite Sequencing. Quality control checks were performed with FastQC v0.11.9 (Brown et al., 2017). Then, Bismark v0.23.1 (Krueger and Andrews, 2011) was used to perform a comparison of the alignments of bisulfite-treated reads to the reference genome downloaded from Phytozome using Bowtie2 v2.3.4.3 with the default parameters (Langmead, 2010). Duplicates were removed via the deduplicate_bismark, and methylation information was extracted by bismark_methylation_extractor.

With the coverage file (*.deduplicated.bismark.cov.gz) as the base, differentially methylated regions (DMRs) were identified using the DSS v2.42.0 R package with smoothing set to true and default parameters (Feng and Wu, 2019). Furthermore, differentially methylated genes (DMGs) were defined as genes with a promoter (2000 kb-long region upstream of mRNA in GTF file) with overlapping DMRs. Un-methylated regions (UMRs) were identified using the MethylseekR v1.34.0 R package (Burger et al., 2013), with parameters m=0.5 and n=4 using BSgenome v1.62.0 package. Furthermore, we identified un-methylated genes (UMG) by the MethylseekR program, which identifies gene promoters that are not considered to be methylated. MethylseekR classifies a segment as a UMR when low methylation levels (<10%) and a relatively high number of CpGs (∼30) are observed (Figure S6).

### miRNA and lncRNA target prediction

The TargetFinder was used to predict the miRNAs target mRNAs (miTmRNAs) and the miRNAs target lncRNAs (miTlncRNAs)(Bo and Wang, 2005). The lncRNAs were predicted to function by regulating the expression of the prospective target genes in a *cis-* or *trans-*acting manner. The protein-coding genes within 100kb of the lncRNA locus were deemed to be targeted in *cis* by the lncRNA (Song et al., 2021). The PCC value was used to analyze the correlations between lncRNAs and mRNAs in samples at 10-time points. The *trans*-acting lncRNAs and their targets were identified with LncTar (Li et al., 2015), with parameter ndG < −0.3.

### Mapman annotation and permutation analysis

The biological function and pathway membership of genes of *B. distachyon* Bd21-3 were annotated using Mercator v4 2.0 (Lohse et al., 2014). The permutation analysis was done as in (Tan et al., 2023). P-values were corrected using the Benjamini-Hochberg correction (Benjamini and Hochberg, 1995).

## Results

### Multi-omic atlas of heat stress acclimation

To identify experimental conditions capturing heat stress memory in *Brachypodium distachyon* Bd21-3, we investigated which temperatures are lethal or can trigger resistance to lethal temperatures (Figure 1A). Heat priming at 39 for one day and 25L for two days of recovery did not negatively impact plant growth (Figure S1). However, when exposed to a heat shock at 42℃ and 48℃ (but not 54℃) for 90 minutes, the primed plants showed less severe yellowing and wilting than the non-acclimated plants (Figure 1B). This suggests that 39L heat priming induces HS memory which can provide at least a 6L tolerance for subsequent HS that occurs two days later. To investigate how the different gene regulatory layers change during the heat priming, we performed a time-course experiment where the non-acclimated (25L acclimated group (39L) were treated with non-lethal HS (39L) after two days (Figure 1C). The first heat treatment has three stages (AA1: 1h before stress, AA2: 3h30m after the start of stress, and AA3: 23h30m after the start of stress), including the recovery phase (AR), while the second heat treatment has the corresponding three stages (AS1-AS3). Samples from all time points were subjected to RNA-Seq, while samples from AA1 (before acclimation), AA3 (at the end of acclimation), and AS1 (before the second heat stress) were also subjected to DNA methylation analysis by bisulfite sequencing and small RNA sequencing by miRNA-Seq (Figure 1C). To control for the differences in plant age between the first and second HS, we also sampled non-acclimated plants at the same age as the second HS (samples NS1-3, Figure 1C).

**Figure 1.**
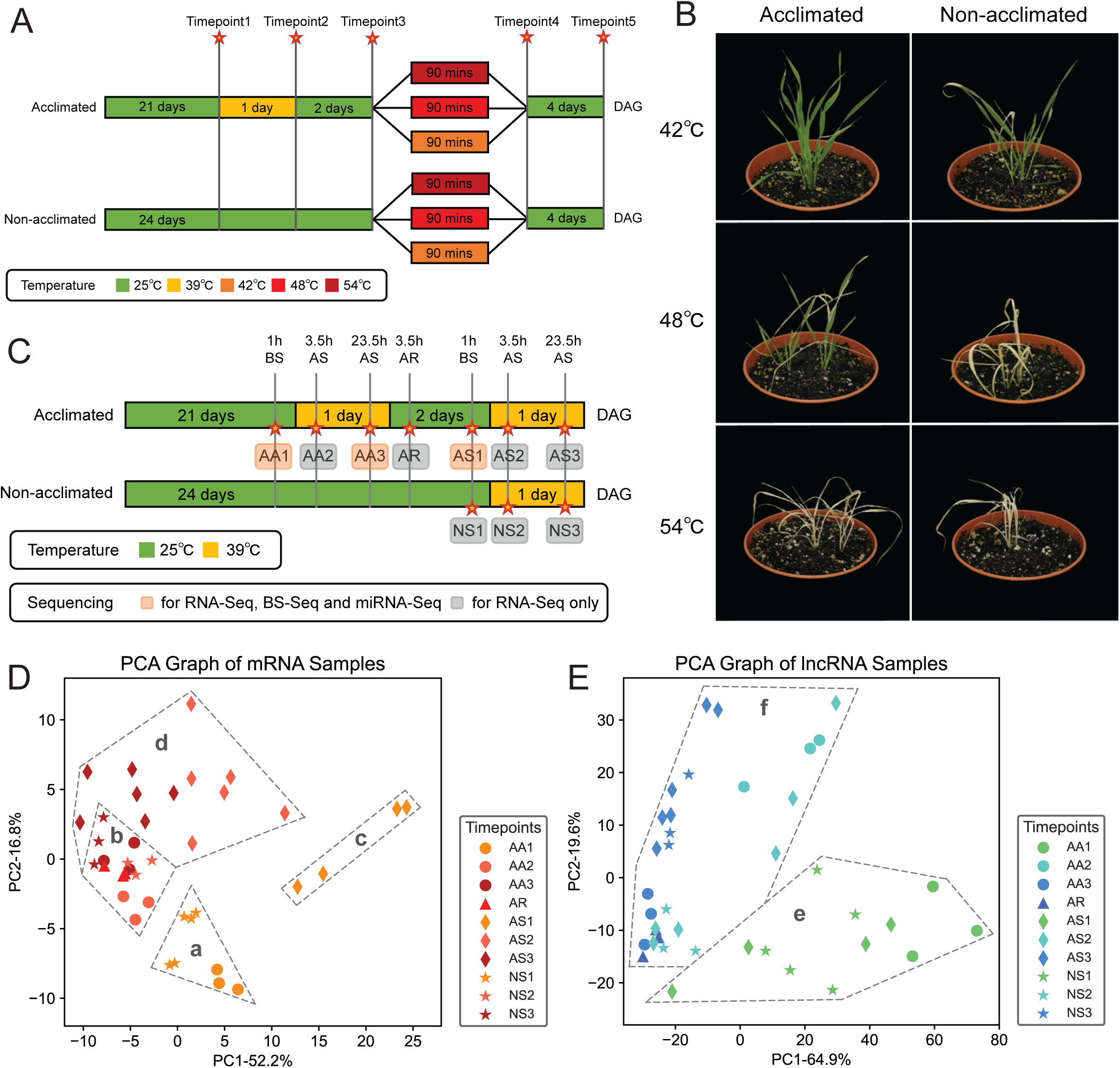
Experimental outline of the heat stress memory experiment in *Brachypodium distachyon*. (A) Heat stress memory experiment. The bar captures days after germination (DAG) with colors representing temperature conditions. The dotted lines and stars point to five distinct imaging time points (Figure S2). (B) Phenotypes of plants four days after heat shock (time point 5). (C) Experimental design for data collection. The dotted lines and stars point to 10 distinct key time points. All time points were analyzed by RNAseq, while time points AA1, AA3, and AS1 were additionally analyzed using BS-Seq and miRNA-Seq. (D) PCA plot of mRNA data. Each point in the PCA graph represents a biological replicate originating from one plant. Different stages are represented by different shapes, with corresponding time points in different stages denoted by different colors. (E) PCA plot of lncRNA data.

Next, we investigated how gene expression changes during the HS experiment (Table S1). The principal component analysis (PCA) of mRNA expression revealed that biological replicates at the same time points were clustered together, indicating the reliability of the sample and sequencing quality (Figure 1D). The time points can be divided into four groups (a, b, c, d) according to their heat stress treatment status - (a) not primed (AA1, NS1), (b) in the heat priming phase or shortly after (AA2, AA3, NS2, NS3, AR), (c) after heat priming and the recovery phase (AS1), and (d) after heat priming and currently in the second HS phase (AS2, AS3).

We also investigated whether long noncoding RNAs (lncRNAs) show similar patterns. To this end, we identified 12,325 lncRNAs (Figure S2) ranging from 200 to 800 nucleotide bases (Figure S3). The PCA plot showed that the samples could be divided into two groups according to the duration of the HS: (e) samples collected before the onset of HS (AA1, AS1, NS1) and (f) samples collected during/after HS (AA2, AA2, NS2, NS3, AR)(Figure 1E). Thus, the PCA of mRNAs expression profiles can differentiate the acclimation status of samples, as, e.g., AA1 is clearly different from AS1. On the other hand, the expression levels of lncRNAs exhibited a similar quality, albeit to a lesser degree as AA1/NS1 are somewhat separated from the acclimated AS1. This suggests that the expression of mRNAs has a closer association with heat stress memory than lncRNAs.

### Gene expression analysis reveals dynamic changes underpinning heat stress acclimation

To understand how gene expression and biological pathways change during heat stress and memory formation, we first performed a k-means clustering analysis to identify distinct expression patterns across all time points.

To identify the optimal number of clusters, we used an elbow plot and inertia analysis, which revealed that 5 clusters provide a good compromise between the number of clusters and inertia (k = 5, Figure S4, Table S2). The five clusters showed distinctive expression profiles (Figure 2A). Out of 50466 transcripts, Cluster 4 and Cluster 5 contained the majority (15900 and 13582), whereas Cluster 1, Cluster 2, and Cluster 3 contained 4778, 5455, and 10751, respectively (Figure 2B). While Clusters 4 and 5 showed either flat or irregular expression profiles with a spike in AS1, respectively, Clusters 1-3 showed more interesting profiles. Clusters 1 and 2 indicate an initial decline in expression after the onset of HS but gradually recover during the heat-duration period, with the highest recovery just before the second HS (Cluster 1, AS1) or earlier, during the recovery phase (Cluster 2, AR). Interestingly, Cluster 3 contains genes that show a continuous increase in expression during HS (AA2, AA3), maintained high expression after the end of the first HS (AR, AS1), and a further increase during the second HS (AS2, AS3). This increasing expression pattern indicates that Cluster 3 might contain genes important for heat stress memory.

**Figure 2.**
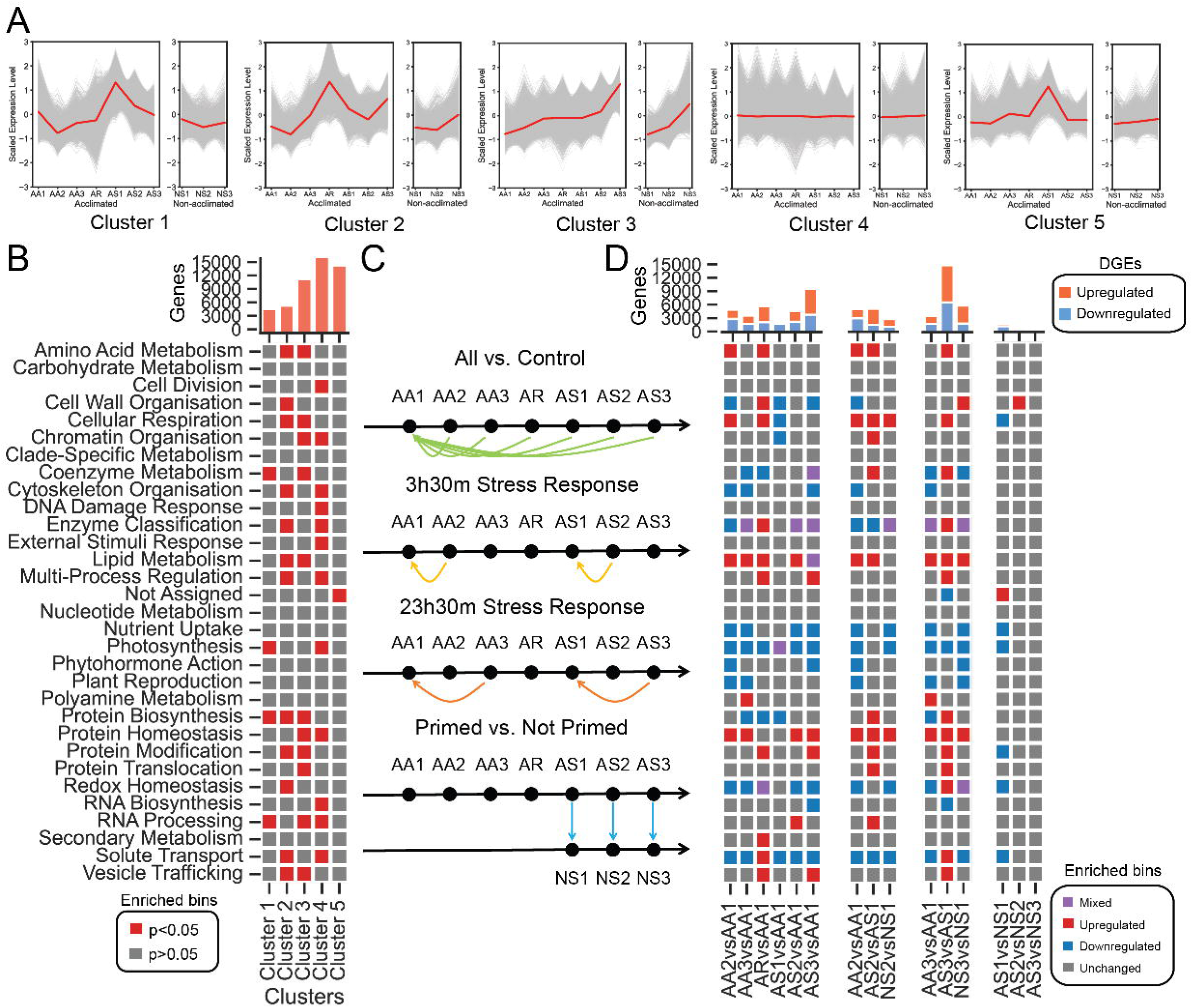
Expression-level signatures during the acquisition of heat stress memory in *Brachypodium distachyon*. (A) Expression patterns of genes in each k-means cluster. In each cluster, the acclimated group and the non-acclimated group were divided into two sub-plots. The z-score normalized expression patterns of individual genes are represented by gray lines, with the average expression patterns of all genes shown with red lines. (B) The bar chart indicates the number of genes found in each k-means clusters. The heatmap indicates the significantly (red) or not significantly (gray) enriched (BH-adjusted p-value < 0.05) Mapman functional categories (rows) for each of the five clusters (columns). (C) Significantly upregulated(red), unchanged (gray), or downregulated (blue) pathways in the four types of comparisons. Purple indicates cases where a significant portion of the genes in a given pathway is down- and upregulated.

To investigate which biological functions are significantly enriched in the five clusters, we performed an enrichment analysis (permutation analysis, Benjamini-Hochberg corrected p-value < 0.05, Table S3). Genes found in Cluster 1 are enriched in photosynthesis and coenzyme metabolism, which agrees with the observation that photosynthesis is decreased during heat stress (Ferrari and Mutwil, 2019)(Figure 2A, red squares). Cluster 2 is enriched for multiple essential processes, such as protein synthesis/modification, respiration, metabolism, and multi-processing regulation, that might be important for signaling during heat stress. Cluster 3, which likely represents genes involved in heat memory, is enriched processes that include many protein-related terms (synthesis, modification, homeostasis, translocation), metabolism (respiration, lipids, coenzymes), and chromatin remodeling. Finally, Clusters 4 (flat) and 5 (irregular) contained various other biological processes.

To obtain a highly detailed view of the changes in gene expression, we identified significantly differentially expressed genes (DEGs) by using different comparison references (Figure 2C, Table S4). For all analyses, we observed thousands of up- and down-regulated genes. Comparing all time points to non-stressed plants (all vs. AA1) revealed downregulation of photosynthesis and phytohormone responses throughout the four days of the experiment and upregulation of protein homeostasis and lipid metabolism (Figure 2C). The short-term responses (3.5 h after the onset of heat: AA2 vs. AA1, AS2 vs. AS1, NS2 vs. NS1) revealed an upregulation of respiration, protein homeostasis, lipid metabolism pathways, and downregulation of photosynthesis, hormone responses, and solute transport. When contrasted with long-term responses (23.5h after the onset of heat: AA3 vs. AA1, AS3 vs. AS1, NS3 vs. NS1), the patterns of differentially expressed biological processes were similar, but cellular respiration genes were no longer differentially expressed. Interestingly, the number of DEGs at the second heat treatment (AS3 vs. AS1) was much higher, indicating that more genes respond to the second stress. The comparison of acclimated vs. non-acclimated plants (AS1-3 vs. NS1-3) showed a comparably low number of DEGs and differentially expressed biological processes (Figure 2D). The highest difference is present before the first heat stress (AS1vsNS1), followed by the short-term responses (AS2vsNS2), and a few differences in long-term responses (AS3vsNS3).

### Heat induces alternative splicing-linked memory

Thermopriming can have a lasting effect on alternative splicing (AS) in Arabidopsis, indicating that alternative splicing can also function as a component in thermomemory (Ling et al., 2018). Therefore, we first set out to investigate the occurrence of the most frequent alternative splicing events: Skipped Exon (SE), Retained Intron (RI), Mutually Exclusive Exons (MXE), Alternative 5S Splice Site (A5SS), and Alternative 3S Splice Site (A3SS). Out of the 50,466 protein-coding transcripts known in the Brachypodium distachyon Bd21-3 v1.2 genome, between 20,000-25,000 showed AS events when compared to unstressed plants (time point AA1, Figure 3A, Table S5). Most of the AS events were of SE and A3SS type, with MXE being the least prevalent (Figure 3A). Next, we identified significantly Differentially Alternatively Spliced Transcripts (DASTs) and observed that the amount of AS events are increased during HS (AA2, AA3, AS2, AS3) when compared to plants in the resting phase (AR, AS1)(Figure 3B, Table S6). Interestingly, the number of DASTs showed a substantial increase during the second heat stress (AS2, AS3 vs. AA2, AA3), which suggests that AS events are also controlled by heat stress memory or might play a role in memory-mediated heat stress tolerance(Figure 3B).

**Figure 3.**
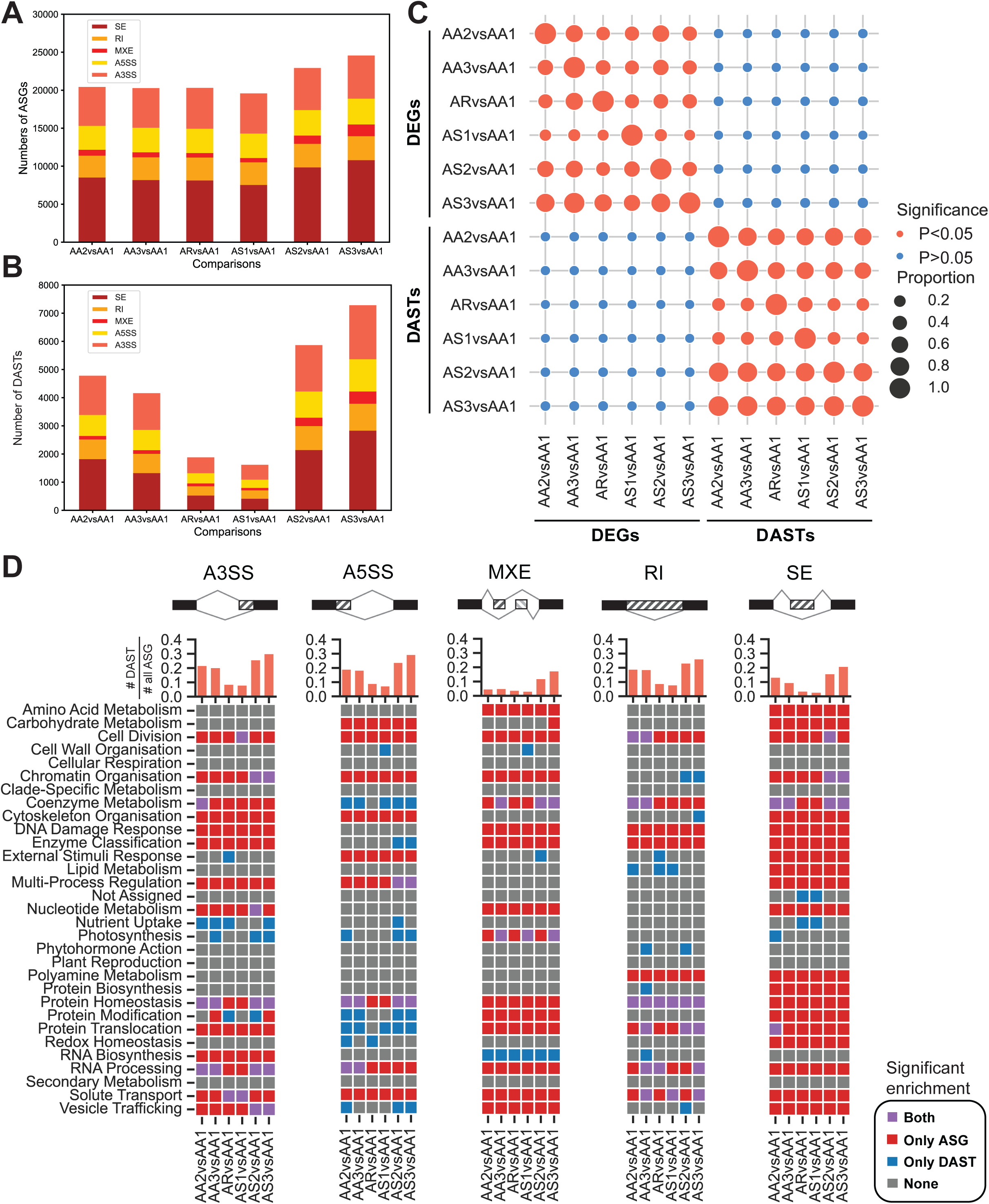
Alternative splicing events during HS. (A) The number and types of alternative splicing events detected for each of the six time points in the heat memory experiment. (B) The number and types of significantly differentially alternatively spliced transcripts (DASTs). (C) The heatmap shows the similarities of DEG and DAST sets between all time points. Red points indicate significant transcript sets (BH-adjusted p ≤ 0.05 identified with a hypergeometric test), while the size of points represents the proportion of the intersection size of two sets (y-axix used as a reference). (D) The proportion and biological processes that DASTs are involved in across the five types of AS events. Bar charts show the proportion of DASTs in each type of AS (number of DASTs/number of AS genes of a given type), while heatmaps display the enriched biological functions of the ASGs (red) and DASTs (blue), or both (purple).

We next set out to investigate whether DEGs also tend to be alternatively spliced by comparing the intersections of DEGs and DASTs at different time points. Interestingly, DEGs and DASTs have no significant intersection (Figure 3C, intersections larger than by chance indicated by red circles). This suggests that DEGs and DASTs are two independent layers regulating mRNAs’ abundance and diversity, respectively.

To elucidate biological processes in which genes undergo alternative splicing, we identified significantly enriched biological pathways to which the AS genes belong (Figure 3D, red cells, BH-adjusted p-value < 0.05, permutation analysis). We observed clear differences between the biological processes and alternative splicing types. For instance, genes involved in photosynthesis are significantly enriched for MXE events throughout all comparisons. SE events tend to have the highest number of genes enriched in various functions, while RI has the lowest. Interestingly, while all AS events show an increase during the second stress, the MXE type shows the most dramatic increase (Figure 3D).

Next, we revealed which biological pathways the DASTs are found in (Table S7). The analysis revealed five patterns of DASTs. 1) Only enriched in the first heat stress (AA2, AA3), indicating a decrease of AS events in the second stress: cell division (RI), coenzyme metabolism (RI), and RNA processing (A5SS). 2) Only enriched in the recovery period (AR, AS1): solute transport (A3SS), nutrient uptake (SE), and not assigned (SE, uncharacterized processes). 3) Only enriched in the second stress (AS2, AS3) indicating AS events influenced by memory: chromatin organization (A3SS, RI, SE), vesicle organization (A3SS), enzyme classification (A5SS) and multiprocess regulation (A5SS). 4) Enriched in both stresses (AA2, AA3, AS2, AS3), indicating AS events that respond to heat: protein homeostasis (A3SS, A5SS), RNA processing (A3SS), protein modification (A5SS) and coenzyme modification (SE). 5) Enriched at all time points, indicating AS events that were sustainably induced by the first stress: RNA biosynthesis (MXE) and protein homeostasis (RI) events.

Taken together, alternative splicing events are highly responsive to HS and might play a major role in establishing HS resilience mechanisms and memory.

### DNA methylation regulates the responsiveness of mRNA levels to heat stress

DNA methylation status is important in the acclimation to stress and memory (Lämke and Bäurle, 2017), as DNA demethylation has been observed in *Arabidopsis thaliana* during heat stress (Korotko et al., 2021). We performed bisulfite sequencing on three key time points (AA1, AA3, AS1) to identify genes with differentially methylated promoters (differentially methylated genes, DMGs). In addition, we also identified genes without significant methylation (unmethylated genes, UMGs, Figure 4A, Table S8).

**Figure 4.**
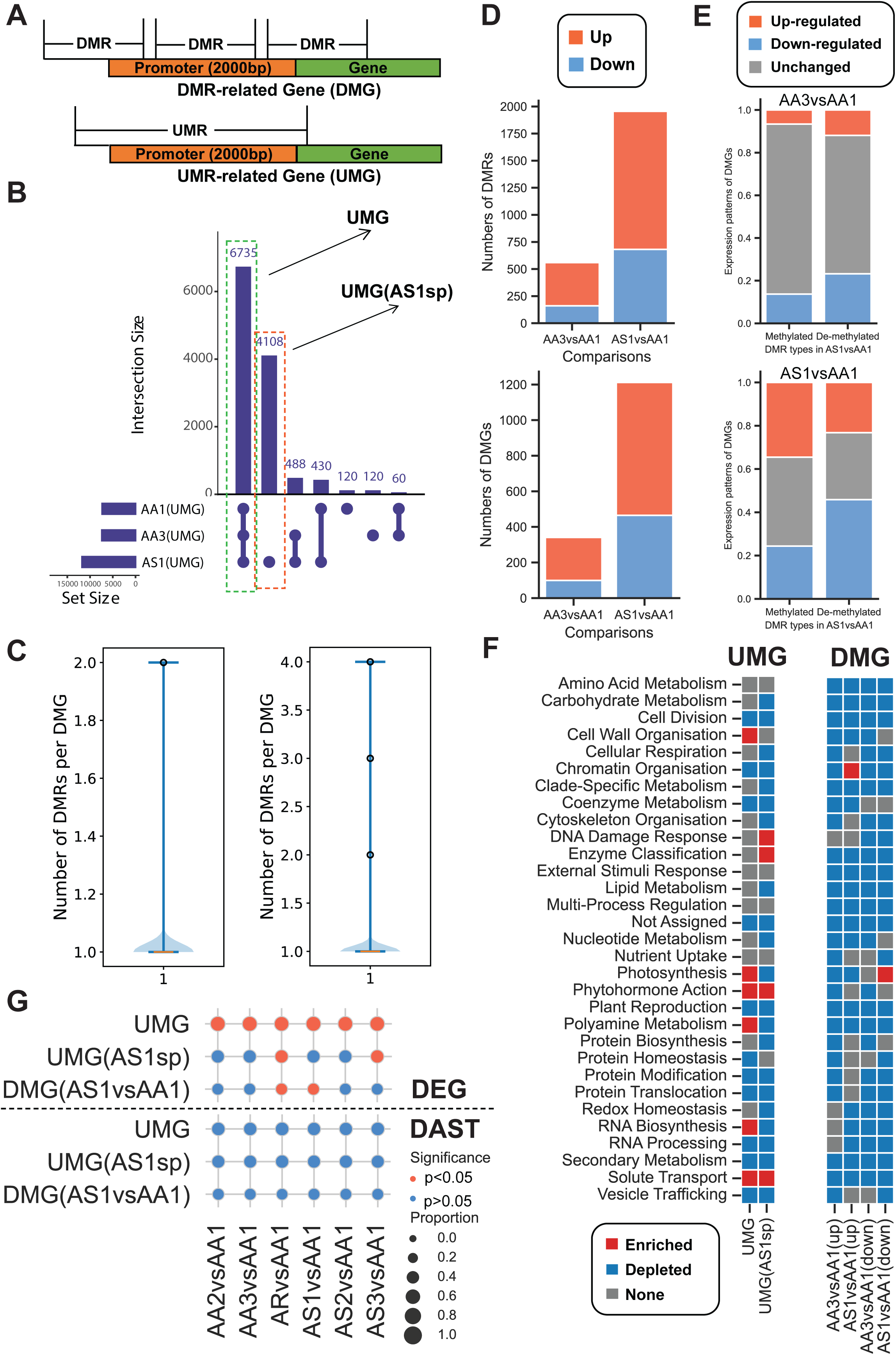
DNA-methylation signatures during heat stress. (A) Definition of differentially methylated region (DMRs) and differentially methylated genes (DMGs), un-methylated regions (UMRs), and un-methylated gene (UMGs). (B) UpsetR plot of UMGs at three time points. The horizontal bar chart depicts the set size of UMGs at the different time points, while the vertical bar chart shows the intersection size among the UMG sets. (C) The number of DMRs associated with DMGs. (D) Number of DMRs and DMGs in AA3vsAA1 and AS1vsAA1 comparisons. (E) Expression correspondence between DMRs and DMGs. Correspondences that pass the hypergeometric test are marked with an asterisk. (F) Biological functions of UMGs and DMGs. Red and blue boxes indicate significantly enriched or depleted biological pathways, respectively (BH-adjusted p ≤ 0.05 identified with a hypergeometric test). Gray boxes indicate no enrichment. (G) The heatmap visualizes the pairwise hypergeometric tests across all UMGs/DMGs and DEGs/DASGs gene sets. In the heatmap, the color of points indicates significant (BH-adjusted p ≤ 0.05 identified with a hypergeometric test) and not significant (p > 0.05) pairs in red and grey, while the size of points represents the proportion of the intersection size of two gene sets to the size of the corresponding gene set on the y-axis.

Most UMGs remained unmethylated at all time points (Figure 4B, 6735 UMG genes in common at AA1, AA3, AS1). However, 4108 new UMGs appeared at AS1, suggesting a large demethylation event after the first heat stress (Figure 4B). Most DMGs have only one DMR, with only a small number having multiple (2 ∼ 4) DMRs that overlap their promoters (Figure 4C, Table S8). For DMGs, we observed a higher number of differentially methylated regions (DMRs) at a later stage (AS1vsAA1) than earlier stage (AA3vsAA1), indicating that DNA methylation can take days to establish after the initial HS (Figure 4D). Interestingly, and in contrast to an increase in unmethylated genes (Figure 4B), most DMGs show an increase in methylation (Figure 4D, orange bars).

Next, we investigated how changes in DNA methylation influences gene expression. To do this, we compared differential gene expression of DMGs where methylation was increased (methylated) or decreased (de-methylated, Figure 4E). Overall, we observed a higher number of DEGs for genes that became de-methylated (Figure 4E, blue bars increase for de-methylated genes). Conversely, genes that became methylated tend not to show changes in gene expression (Figure 4E, gray bars).

Next, we investigated the biological pathways to which the UMGs and DMGs belong (Table S9). UMGs are typically underrepresented (i.e., fewer UMGs in a pathway than expected by chance, permutation analysis, BH-adjusted p-value < 0.05), but few pathways such as cell wall organization, photosynthesis, and phytohormone action contain more UMGs than expected (Figure 4F, red cells, UMGs). The pathways for UMGs that appeared after the first stress (UMG AS1sp) included DNA damage response and enzyme classification (Figure 4F). In contrast, the biological functions of the observed DMGs (Figure 4F) were unclear, as most biological pathways were not enriched (Figure 4D).

Finally, we investigated the relationships (if any) that methylation shares with the expression level and alternative splicing of genes. We observed a significant overlap between sets of DEGs and UMGs (Figure 4G) across all time points (Figure 4G, BH-adjusted p-value < 0.05, hypergeometric test). This suggests that unmethylated genes are more likely to change their expression than genes with methylated promoters. Conversely, there is a lack of evidence indicating that DNA methylation plays a role in the system-wide regulation of alternative splicing within the context of heat stress and memory, as none of the differentially alternatively spliced genes showed a significant overlap with UMGs or DMGs (Figure 4G).

To conclude, we observed that the first heat stress led to DNA demethylation (Figure 4B) and a higher number of differentially methylated genes (Figure 4D). Furthermore, unmethylated genes tend to show changes in gene expression (Figure 4G), suggesting that promoter demethylation plays a significant role in the transcriptional regulation of DEGs at the second heat stress.

### Long noncoding RNAs can influence mRNA levels

Long noncoding RNAs (lncRNAs) are riboregulators that can control gene expression at the transcriptional, post-transcriptional, and epigenetic levels by targeting various stress-responsive mRNAs, transcription factors, and microRNAs (Jha et al., 2020). While the involvement of lncRNAs in plants’ response and memory formation during HS has yet to be experimentally validated, various studies have shown that lncRNA are highly responsive to heat, suggesting their potential involvement in abiotic stress survival (Xin et al., 2011; Wang et al., 2019).

To study the role of 12325 lncRNAs (Table S10) in HS, we first identified the putative mRNA targets of the 12325 lncRNAs (see methods, Table S11). Overall, we found that *cis-* and *trans-*acting lncRNAs tend to target a similar number of genes (median ∼20 for both, Figure 5A), but *trans-*lncRNAs can have a larger number of mRNA targets (Figure 5A). Conversely, one gene tends to have one and, at most, two *cis-*lncRNAs targeting it, while up to eight *trans-*lncRNAs can target a given mRNA (Figure 5A). In total, 483 lncRNAs showed differential expression (Table S12), with the majority showing significant upregulation at the last time point of the second heat stress (Figure 5B, AS3, orange bar).

**Figure 5.**
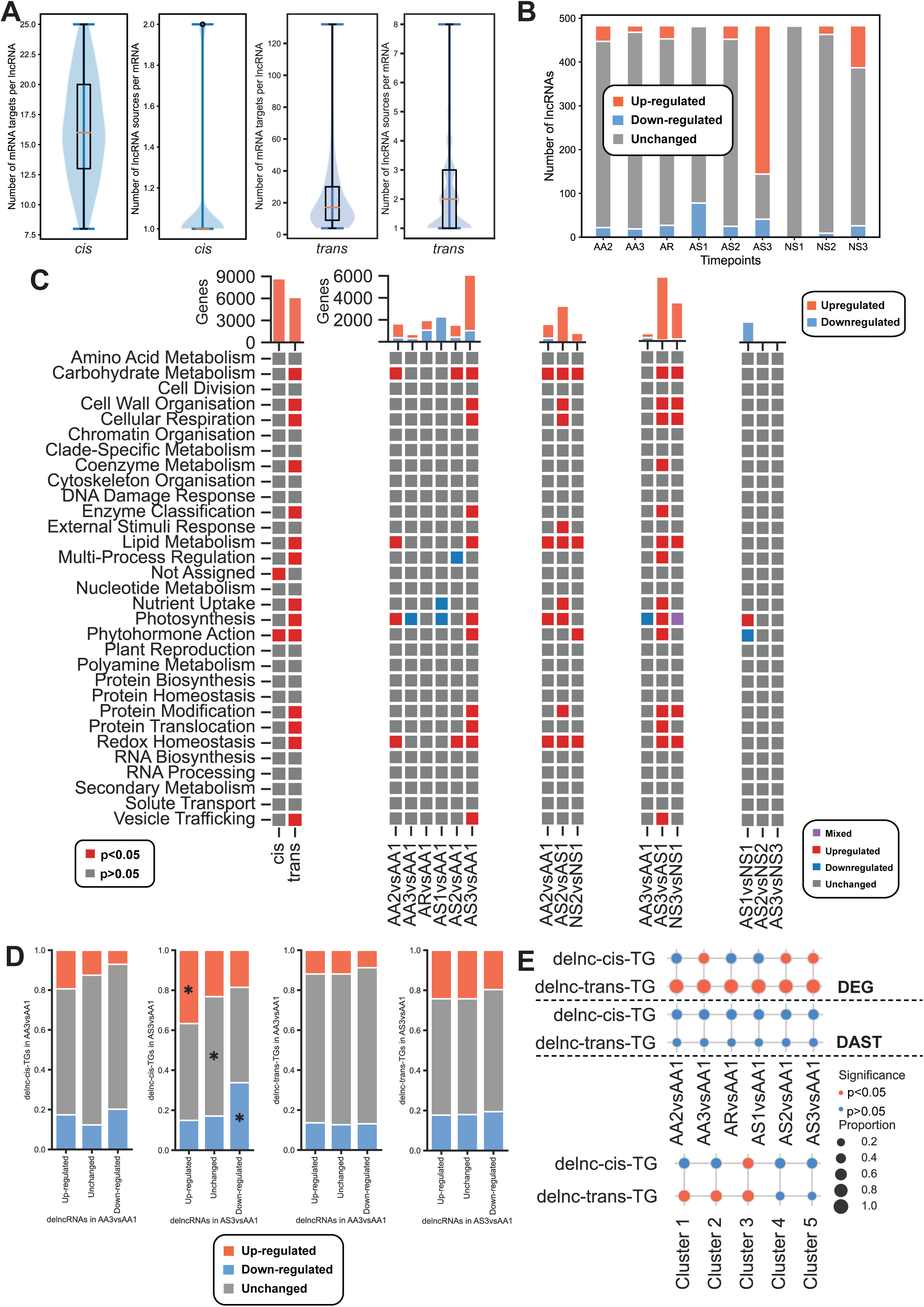
lncRNA signatures during HS. (A) The number of mRNA targets per lncRNA and the number of lncRNAs per mRNA for *cis-* and *trans-*acting lncRNAs. (B) The number of differentially expressed lncRNAs across the time points. Orange, blue, and gray bars indicate up-, down-regulated, or unchanged expression, respectively. (C) Biological functions of the differentially expressed *cis-* and *trans-*acting lncRNA. Bar charts illustrate the number of genes targeted by each lncRNA class. Biological functions of differentially expressed lncRNAs at different time points. Red and blue boxes indicate significantly enriched or depleted biological pathways, respectively (BH-adjusted p ≤ 0.05 identified with a hypergeometric test). Gray boxes indicate no significant enrichment. Bar charts illustrate the number of genes targeted by each lncRNA class. Red and blue bars in the bar charts represent up- and down-regulated genes. (D) Expression correspondence between lncRNAs and mRNAs, where orange, gray, and blue bars indicate up-, unchanged, and down-regulated mRNA targets. Correspondences that are larger than expected by chance are marked with an asterisk. (E) Pairwise hypergeometric tests testing for the enrichment of the lncRNA gene targets, including DEGs, DAST, and k-means clusters. The analysis was performed as in Figure 4G.

We performed a functional enrichment of their mRNAs targets to elucidate the biological processes targeted by the lncRNAs (Figure 5C). While mRNA targets of *cis-* lncRNAs (8696 targets) are only enriched in phytohormone responses and not assigned category (indicating poorly characterized biological processes), *trans-*lncRNAs (6140 targets) belong to functional categories such as lipid metabolism, multiprocess regulation, protein-related processes, and others (Figure 5C), which are similar processes that we observed for DEGs (Figure 2B-C).

Next, we investigated how lncRNAs can influence the expression of the target genes. To do this, we tested the correspondence between three possible states (up-, down-regulated, unchanged) of lncRNAs and their predicted mRNA targets. Only for *cis-*acting lncRNAs a significant association was found, where upregulated *cis-* ting lncRNAs (x-axis) are significantly associated with up-regulation of their targets (y-axis, Figure 5D, orange bar, the asterisk indicates BH-adjusted p ≤ 0.05). Furthermore, the down-regulation of *cis-*lncRNAs was also associated with a significant decrease in their mRNA targets (Figure 5D, asterisk, blue bar). However, no significant associations were found for trans-acting lncRNAs, indicating that lncRNAs either do not change the expression of their targets or that the change could be both positive or negative.

Finally, we investigated whether differentially expressed lncRNAs are associated with the DEGs at the different time points and the five k-means clusters. We first tested whether the mRNA targets of differentially regulated lncRNAs are more present than expected by chance in the observed DEGs. We observed that DEGs are enriched for *cis-* (enriched at time points AA3, AS2, AS3) and *trans-*acting (all time points) lncRNAs (Figure 5D). However, the targets of lncRNAs are not enriched for genes that undergo alternative splicing, suggesting that lncRNAs do not influence splicing (Figure 5E, DAST). The targets of the lncRNAs are also enriched for genes found in k-means clusters that show responses to heat treatments (clusters 1, 2, and 3), but not 4 (flat) or 5 (irregular), of which cluster 3 (likely involved in heat stress memory) is significantly targeted by both *cis-* and *trans-*acting lncRNAs.

### Small RNAs are associated with alternative splicing

Small RNAs negatively regulate gene expression by directing target mRNA cleavage, translational repression, and DNA methylation (Li et al., 2017a). In addition, small RNAs have been implicated in controlling phenotypic plasticity, abiotic/biotic responses, and symbiotic/parasitic interactions (Song et al., 2019). There is evidence for miRNAs regulating the heat stress response in various plants (Guan et al., 2013; Stief et al., 2014; Ding et al., 2017), but the roles of miRNAs in Brachypodium are unclear.

To identify the miRNAs in our dataset, we downloaded all available, mature 48885 miRNAs (size 15-34 nt) from mirBase (Table S13), of which 34683 had a non-zero expression in our dataset. The majority of miRNAs had one predicted mRNA target (median two targets), and similarly, most targeted mRNAs were targeted by only one miRNA (median four miRNAs)(Figure 6A, Table S14). LncRNAs can also be targeted by miRNAs and function as a molecular ‘sponge’ to protect the intended targets (Meng et al., 2021). We similarly identified a predominantly 1-to-1 correspondence of miRNA to lncRNA (Figure 6A).

**Figure 6.**
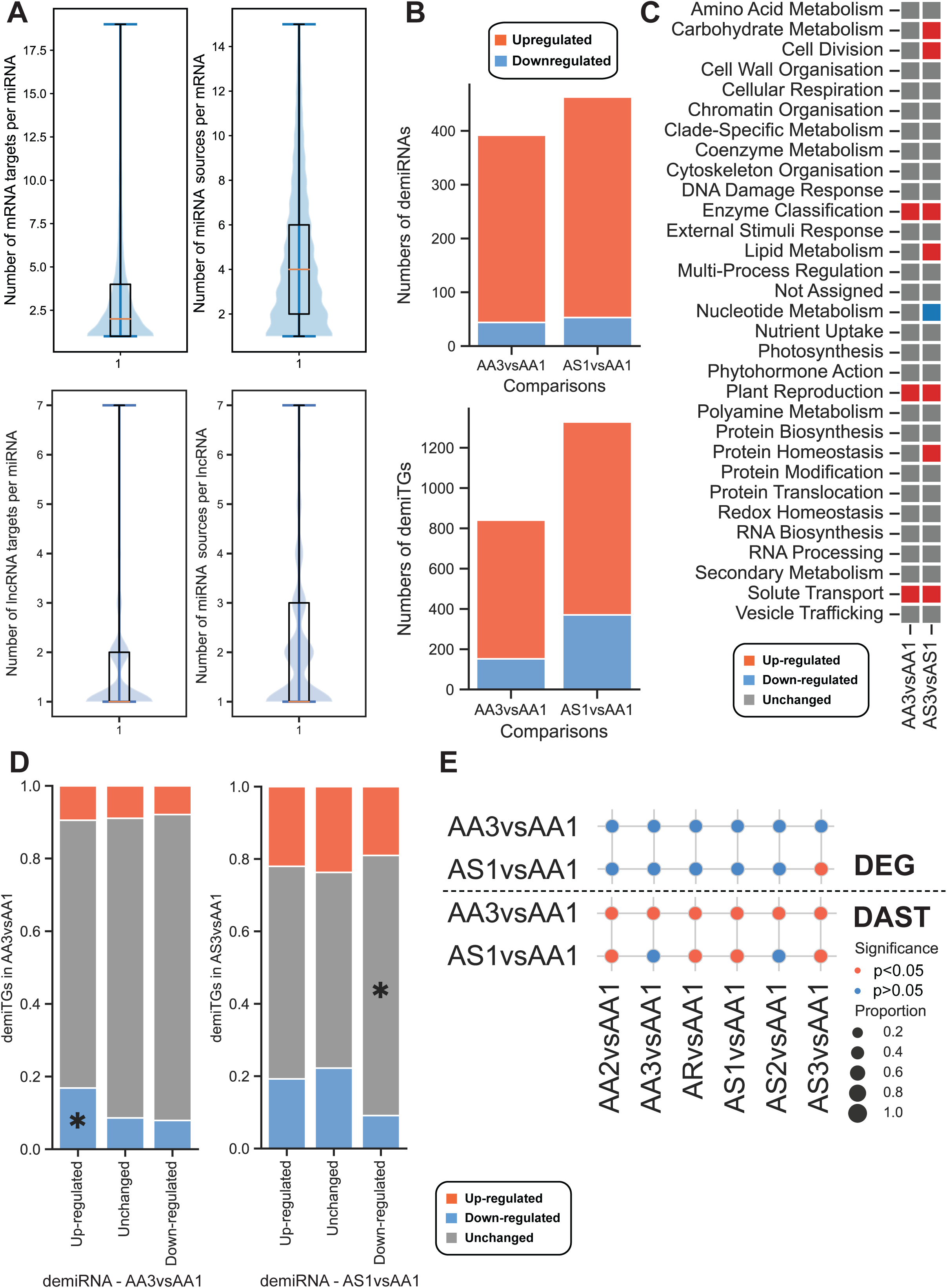
miRNA-level signatures during heat stress. (A) The number of mRNA targets per miRNAs and miRNA per mRNA. (B) The number of significantly differentially expressed miRNAs (demiRNAs) and differentially expressed miRNAs target genes (demiTGs) at AA3vsAA1 and AS1vsAA1. (C) Biological functions of demiTGs. Red and blue boxes indicate significantly enriched or depleted biological pathways, respectively (BH-adjusted p ≤ 0.05 identified with a hypergeometric test). (D) Expression correspondence between miRNAs and mRNAs, where orange, gray, and blue bars indicate up-, unchanged, and down-regulated mRNA targets. Correspondences that are larger than expected by chance are marked with an asterisk. (E) The heatmap visualizes the pairwise hypergeometric tests testing for the enrichment of the miRNA gene targets, including DEGs and DASTs. The analysis was performed as in the previous figures.

Next, to better understand how heat stress and memory influence the expression of miRNAs, we identified differentially expressed miRNAs (demiRNAs) at the two critical time points AA3 and AS1 (Table S15). Most demiRNAs were upregulated at both time points, with a higher number of demiRNAs at the later time point (Figure 6B). Surprisingly, by analyzing the miRNA-target pairs, we observed that the predicted target genes (demiTGs) of demiRNAs are also predominantly upregulated (Figure 6B), which is not in line with the expected negative correlation between miRNA and the target. The biological functions of the targeted genes include various enzymatic functions (enzyme classification), solute transport, and plant reproduction (Table S16), and also protein homeostasis, lipid and carbohydrate metabolism, and cell division at the later time point (Figure 6C).

To better understand how demiRNAs can influence the expression of their target genes (demiTGs), we performed a similar analysis as for DNA methylation and lncRNAs. We investigated the correspondence between three possible states (up-, down-regulated, unchanged) of demiRNAs and their demiTGs. Overall, we observed that up-regulation of miRNAs is associated with the downregulation of DEGs at the early time point (AA3, Figure 6D, the asterisk indicates BH-adjusted p ≤ 0.05), but not at the later time point (AS1). However, overall, we observed only a minor association between the differential expression of miRNAs and their targets across the time points (Figure 6E, DEG analysis, only AS3 is significant), suggesting that the miRNAs play only a modest role in controlling mRNA levels in our experiment. Surprisingly, we observed a significant association between demiTGs and DASTs, indicating that the targets of miRNAs tend to show alternative splicing (Figure 6E).

To summarize, many miRNAs are upregulated due to heat stress (Figure 6B) and are associated with the alternative splicing of their target genes (Figure 6E).

### Comparative, multi-omic dissection of heat stress memory in flowering plants

Gene expression patterns that are conserved across species are likely to be enriched for relevant biological functions (Hansen et al., 2014). Since gene expression patterns underlying heat stress memory have been extensively studied in Arabidopsis thaliana (Oberkofler et al., 2021), we set to compare the conservation of transcriptomic responses between Arabidopsis and Brachypodium. However, different abiotic stresses typically induce an unspecific kingdom-wide transcriptional program that results in the downregulation of several pathways, including photosynthesis (Ferrari and Mutwil, 2019), as we have also observed for Brachypodium (Figure 2D). Since these downregulation patterns are not specific to any given stress, and we aim to understand better which genes and pathways are activated to mediate stress resilience and memory, we focused on identifying conserved, upregulated gene expression responses. The conserved responses are thus likely capturing heat memory responses in the ancestor of flowering plants.

To identify the conserved responses, we first obtained gene expression data from an HS experiment from Arabidopsis (Liu et al., 2018)(Table S17). The dataset captures three variations of heat treatment: P (primed), T (triggered), P+T (primed then triggered), and a non-treated control N (naive)(Figure 7A). Then, to compare the transcriptomic responses across species, we identified the upregulated gene families in Arabidopsis and Brachypodium and calculated whether the sets of upregulated gene families were more similar than expected by chance. The transcripts that are upregulated in Arabidopsis comprise HS-responsive genes (upregulated in T vs. N), sustained upregulation after HS (P vs. N, i.e., type I memory) (Oberkofler et al., 2021), and enhanced re-induction upon a recurrent HS (P+T vs. P, type II memory). We performed an exhaustive comparison of all combinations, time points, and clusters. We observed that all time points in Brachypodium are significantly similar to the HS responses in Arabidopsis (Figure 7B, T vs. N, BH-adjusted p ≤ 0.05). Furthermore, the sustained genes in Arabidopsis (P vs. N) were significantly similar to clusters 2 and 3, and genes upregulated during the second HS (AS2vsAS1, AS3vsAS1, AS1vsAA). Similarly, Arabidopsis genes that showed enhanced (P+T vs. P) responses were also similar to clusters 2 and 3 and HS responses in Brachypodium (Figure 7B). Interestingly, Brachypodium cluster 3, likely containing genes important for HS response and memory, is significantly similar to nearly all heat stress treatments in Arabidopsis (Figure 7B). Thus, we conclude that the HS responses and likely memory mechanisms are conserved between monocots and dicots.

**Figure 7.**
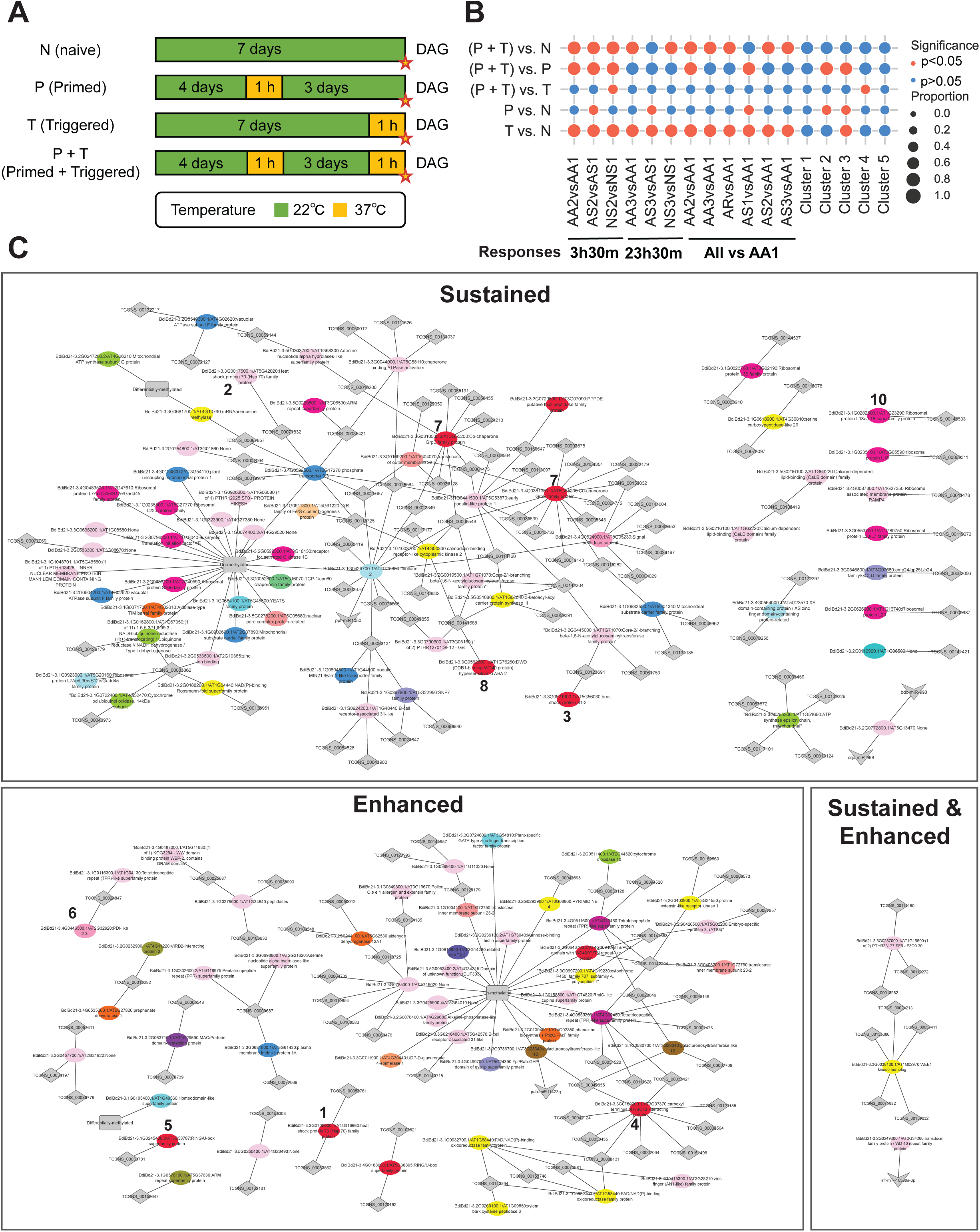
Comparative heat stress memory analysis between *Brachypodium distachyon* and *Arabidopsis thaliana*. (A) Design of the heat stress memory experiment in *Arabidopsis thaliana*. The bar illustrates the timeline of the day after germination (DAG), the different colors represent various temperature conditions, and the stars indicate sample collection time points. The four groups are: N (naive, no HS), P (primed, 1 hour heat priming at 4DAG), T (triggered, 1 hour heat at 8 DAG), and P+T (1 hour heat at 4 and 8 DAG). (B) The heatmap visualizes the pairwise hypergeometric tests testing for the enrichment of the Brachypodium genes upregulated at the different time points and clusters (x-axis) and the Arabidopsis genes upregulated at the different combinations of N, P, T, and P+T. The statistical analysis was performed as in the previous figures. (C) Conserved genes and their regulators are indicated by dark and light shades, respectively. The regulators from different layers are depicted in different shapes: DNA methylation (rectangles), lncRNAs (diamonds), and miRNAs (arrows). The node colors depict different Mapman functional categories, where genes indicated in protein quality control and biosynthesis are indicated by red an purple colors, respectively.

To identify these conserved HS responses that were likely found in the ancestor of flowering plants, we identified the conserved sustained and enhanced genes of Arabidopsis and genes found in cluster 3 in Brachypodium (Table S18). We also plotted the noncoding RNAs and DNA methylation states that likely regulate these genes’ expression in Brachypodium (Figure 7C). The network revealed several known genes involved in heat stress tolerance and memory (red colors), such as: Heat Shock Proteins 70 (BdiBd21-3.3G0702600, indicated by 1 in the enhanced cluster), (BdiBd21-3.3G0017500, 2, sustained cluster) and HSP 81-2 (BdiBd21-3.3G0521900.1, indicated by 3, sustained), ubiquitin ligases important for HS tolerance (BdiBd21-3.3G0180200.1, 4, enhanced)(Zhou et al., 2014), a ubiquitin ligase essential for stress hormone abscisic acid response (BdiBd21-3.1G0245400.2, 5, sustained)(Zhou et al., 2014), disulfide isomerase necessary for pollen survival during HS (BdiBd21-3.4G0446500.1, 6, enhanced)(Feldeverd et al., 2020), nucleotide exchange factor of the HSP70 complex (BdiBd21-3.4G0381300.1, 7, sustained)(Hu et al., 2012) and DDB1-binding WD40 proteins that are hypersensitive to ABA (BdiBd21-3.3G0562800.1, 8, sustained)(Lee et al., 2010). Other genes that are conserved between cluster 3 and the Arabidopsis HS memory genes include plastid metalloprotease important during HS memory (Sedaghatmehr et al., 2016), and E3 ligases involved in ABA responses during drought (Table S18)(Yang et al., 2018). The network also contained numerous ribosomal proteins (10, nodes indicated by purple color), which aligns with their proposed role in acclimation to stress (Garcia-Molina et al., 2020).

Interestingly, many of the abovementioned genes tend to be connected to one or more lncRNAs (Figure 7C, indicated by diamonds), as, e.g., the nucleotide exchange factor (BdiBd21-3.4G0381300.1, 7, sustained) is potentially regulated by ten lncRNAs (Figure 7C). Most of the identified regulatory interactions for the discussed proteins involve one or more lncRNAs, suggesting their role in regulating the expression of genes involved in protein homeostasis and biosynthesis.

Taken together, we showed that the HS memory responses are conserved between Arabidopsis and Brachypodium, suggesting that lncRNAs are important regulators of these genes.

## Discussion

The heat stress that plants experience due to global warming significantly impacts both their growth and the production of crops. Understanding how plants react, acclimatize to, and survive HS has become increasingly crucial as the risk of crop loss and the need for food worldwide continue to rise. As extreme high temperature episodes in the natural environment are frequently clustered in time, HS memory is an adaptation mechanism that enables plants to respond more quickly to recurring HS events. In Arabidopsis, primed plants can better cope with HS up to five days after priming (Charng et al., 2006; Stief et al., 2014). Numerous molecular mechanisms underpinning HS memory have been reported. Since *Brachypodium distachyon* is a model plant for grasses and no HS memory studies have been performed in this species, we set to explore the temporal changes of gene expression, alternative splicing, (non)-coding RNAs, and DNA methylation across two HS events. We observed that Brachypodium could acclimatize to HS as the plant showed healthier phenotypes at 48*C when compared to a naive plant (Figure 1B).

Transcripts under the control of HS memory show Type I: sustained induction after HS for several days (Stief et al., 2014) and/or Type II: enhanced re-induction upon a recurrent HS event (Lämke et al., 2016). In line with previous studies, a second exposure to HS resulted in a unique mRNA expression pattern (Figure 1D) with a higher number of differentially expressed genes in Brachypodium (Figure 2C). Clustering analysis of expression profiles revealed five clusters, of which cluster 3 showed a continuous increase of gene expression during HS, thus capturing genes with a mixture of Type I (sustained) and Type II (enhanced) expression patterns (Figure 2A). The biological function of the genes in cluster 3 is also in line with the biological processes important for HS survival, acclimation, and memory: protein (synthesis, modification, homeostasis, translocation)(Zhou et al., 2014; Garcia-Molina et al., 2020), metabolism (respiration, lipids)(Shiva et al., 2020; Scafaro et al., 2021) and chromatin organization (Huang et al., 2023).

In line with the previous studies on alternative splicing and HS (Ling et al., 2018), we observed that AS also shows Type I and Type II memory patterns (Figure 3B). Splicing changes under HS have been seen in different eukaryotic systems and may thus represent a mechanism shared among eukaryotic species (Chang et al., 2014). However, in contrast to a study in Arabidopsis that observed retained introns (RI) as the major form of AS (Ling et al., 2018), we observed skipped exons (SE) as the predominant AS event (Figure 3A, B). Furthermore, while the study in Arabidopsis observed fewer IR events during HS of primed plants (Ling et al., 2018), we observed the opposite, as a substantially higher number of transcripts showed AS across all categories (Figure 3B). This suggests that AS responses might be species-specific, and more studies are needed to identify conserved trends. Surprisingly, we found no significant overlap between differential gene expression and differential splicing (Figure 3C), suggesting that gene expression and AS are not linked. Interestingly, the various types of AS events showed preferences for diverse biological processes (Figure 3D), where most biological pathways were enriched in at least one type of AS event at some point. This highlights that AS might have a dramatic, underappreciated role in HS survival and memory and merits further study in more than one species.

Genome-wide DNA demethylation has been observed in multiple species after HS (Min et al., 2014; Hossain et al., 2017; Korotko et al., 2021), which aligns with an increase in unmethylated genes in our study (Figure 4B). While the exact role of DNA methylation in the context of HS survival and memory is unclear (Liu and He, 2020), we observed a significant overlap between genes that are differentially expressed (with no link to alternative splicing) and un-methylated (Figure 4G), suggesting that demethylation could boost ‘flexibility’ of gene expression after the initial HS. While DNA methylation is typically linked with the down-regulation of gene expression (Zilberman et al., 2007), we observed that un-methylated genes are significantly downregulated (Figure 4G), further suggesting that demethylation mediates increased variability of gene expression after HS.

The function of lncRNAs in HS survival and memory is unclear (Zhao et al., 2020), despite a study reporting that lncRNAs are responsive to HS (Song et al., 2016). LncRNAs are thought to indirectly regulate gene expression by competing for binding regulatory RNAs (e.g., miRNAs)(He et al., 2020). We identified 12325 lncRNAs based on RNA-seq data, of which 483 showed differential expression. We predicted a large cis- and trans-acting lncRNA-mRNA regulatory network (Table S14), which revealed an association between lncRNAs and differential gene expression (Figure 5E). In contrast, the identified miRNAs and the analysis of the inferred miRNA-mRNA regulatory network revealed a weak association between miRNA targets and differential gene expression but a significant link with alternative splicing of the target genes (Figure 6E).

To uncover flowering plant ancestral genes that are important for HS survival and memory, we identified genes that show a conserved sustained and enhanced expression in Arabidopsis and Brachypodium. The identified genes comprised many known genes important in protein homeostasis and biosynthesis (Figure 7C) but also revealed many genes without any known function (Table S18). Since the identified genes show conserved type I and II HS responses between Arabidopsis and Brachypodium, these genes comprise prime candidates for future study. Interestingly, we observed many associations between the conserved genes and lncRNAs, which warrants further investigation.

The HS response mechanism of genes of model plants is beginning to take shape after years of research (Zhao et al., 2020). However, the functions of the different gene regulatory layers and their interconnections are still poorly understood. In this study, we revealed several unknown interconnections. We highlighted several differences between the responses of Arabidopsis and Brachypodium, highlighting that more studies are needed at the different regulatory layers in more species. Furthermore, as all studied layers have shown evidence of HS memory, it is likely that many yet unknown mechanisms are also active. Other epigenetic modifications, such as phosphorylation, ubiquitination, and SUMOylation (Small Ubiquitin-like Modifier, SUMO), RNA methylation, cell type-specific expression (studied by, e.g., single-cell RNAseq), need to be studied to more comprehensively dissect the composition and interconnectivity of the molecular mechanisms underpinning HS survival and memory.

## Data availability

The RNA-seq, sRNA-seq, and mDNA-seq data are available at E-MTAB-12755, E-MTAB-12739, and E-MTAB-12727, respectively.

## Acknowledgments

A China Scholarship Council fellowship sponsors XZ. MM is funded by the Ministry of Education Tier 2 grant MOE-T2EP30122-0017.

## Author contributions

XZ extracted the DNA and RNA, sent samples for sequencing, and performed data analysis. QWT coordinated the HS experiments and provided suggestions for the data analysis. RP performed the HS experiment. PKL co-supervised XZ and provided suggestions for the data analysis. MM conceived the project with help from QWT, acquired funding, provided suggestions for the data analysis, and co-wrote the paper with XZ. The authors thank all the members of the Mutwil Lab for their feedback during lab meetings.

---

**Table S1. TPM values of transcripts**. The columns represent different time points, while rows capture transcripts.

**Table S2. Transcript to k-means cluster.** The columns represent different clusters, while rows capture transcripts. Values of 1 and 0 indicate cases where a transcript does/does not belong to the cluster, respectively.

**Table S3. Mapman analysis of clusters.** The columns represent the clusters and the different comparisons, while the rows represent the Mapman bins. The numbers indicate the adjusted p-values.

**Table S4. DGE analysis of mRNAs.** The columns represent the different comparisons, while the rows represent transcripts. Values of 1 and 0 indicate changed (up or down) and unchanged transcript levels.

**Table S5. Alternative splicing events during the HS experiment.** The numbers indicate transcripts (rows), while columns capture the different time points. Values of 1 and 0 indicate cases where a transcript does/does not show AS, respectively.

**Table S6. DGE of alternative splicing.** The numbers indicate transcripts (rows), while columns capture the different time points. Values of 1 and 0 indicate cases where a transcript is/is not a DAST, respectively.

**Table S7. Mapman analysis of DASTs.** The columns represent the different comparisons, while the rows represent the Mapman bins. The numbers indicate the adjusted p-values.

**Table S8. DNA methylation occurrences and adjusted p-values at the different time points (columns).** The rows capture transcripts. Values of 1 and 0 indicate changed (up or down) and unchanged DNA methylation levels of the affected transcripts.

**Table S9. Mapman analysis of DNA methylation.** The columns represent the different comparisons, while the rows represent the Mapman bins. The numbers indicate the adjusted p-values.

Table S10. Sequences of lncRNAs.

**Table S11. Targets and types of lncRNAs.** The columns indicate the source and targets of the lncRNAs.

**Table S12. DGE of lncRNAs.** Values of 1, 0, −1 indicate up-, unchanged, and down-regulated lncRNAs.

Table S13. Sequences of miRNAs

**Table S14. Table with miRNA-mRNA, miRNA-lncRNA pairs.** The columns indicate the source and targets of the miRNAs.

**Table S15. DGE of miRNAs.** Values of 1 and 0 indicate changed (up or down) and unchanged miRNA levels.

**Table S16. Mapman analysis of miRNA targets.** The columns represent the different comparisons, while the rows represent the Mapman bins.

**Table S17. DGE analysis of *Arabidopsis* genes.** The columns capture *Arabidopsis* probesets (1st column), DGE responses (1, 0, −1 up-, unchanged, down-regulated) of the different comparisons (2nd-6th column), and the best hit homolog of *Brachypodium* (column H) and the *Arabidopsis* gene (I).

**Table S18. Conserved HS memory network**. Brachypodium transcripts (column 1), any regulatory molecules/modifications (2), memory type of Arabidopsis best blast hits (3), membership of Brachypodium cluster (4), and the type of the source node.

Figure S1. The growth phenotypes of *Brachypodium* plants subjected to the different HS treatments.

Figure S2. The pipeline used to identify the lncRNAs.

Figure S3. The length distribution of the identified lncRNAs

Figure S4. Identification of the k value. The elbow point, the percentage of explained variance with the different PCs, and the PCA plots of the different PC combinations are shown.

Figure S5. UpSetR plot analysis of the alternatively spliced genes and differentially alternatively spliced transcripts. The plots show the similarities and differences between the sets.

**Figure S6. MethylSeekR analysis of the CpGs feature of the segments on the genome level.** The plots show the number of CpGs (hypomethylated regions) versus their average methylation level for the different time points, in triplicates.

## References

Asensi-Fabado, M. A., Amtmann, A., and Perrella, G. (2017). Plant responses to abiotic stress: The chromatin context of transcriptional regulation.

Battisti, D. S., and Naylor, R. L. (2009). Historical warnings of future food insecurity with unprecedented seasonal heat. Science 323:240–244.

Benjamini, Y., and Hochberg, Y. (1995). Controlling the False Discovery Rate: A Practical and Powerful Approach to Multiple Testing. Journal of the Royal Statistical Society: Series B (Methodological) 57:289–300.

Bo, X., and Wang, S. (2005). TargetFinder: a software for antisense oligonucleotide target site selection based on MAST and secondary structures of target mRNA. Bioinformatics 21:1401–1402.

Bolger, A. M., Lohse, M., and Usadel, B. (2014). Trimmomatic: A flexible trimmer for Illumina sequence data. Bioinformatics 30:2114–2120.

Boyko, A., Blevins, T., Yao, Y., Golubov, A., Bilichak, A., Ilnytskyy, Y., Hollander, J., Jr, F. M., and Kovalchuk, I. (2010). Transgenerational Adaptation of Arabidopsis to Stress Requires DNA Methylation and the Function of Dicer-Like Proteins. PLOS ONE 5:e9514.

Brown, J., Pirrung, M., and McCue, L. A. (2017). FQC Dashboard: integrates FastQC results into a web-based, interactive, and extensible FASTQ quality control tool. Bioinformatics 33:3137–3139.

Brzezinka, K., Altmann, S., Czesnick, H., Nicolas, P., Gorka, M., Benke, E., Kabelitz, T., Jähne, F., Graf, A., Kappel, C., et al. (2016). Arabidopsis FORGETTER1 mediates stress-induced chromatin memory through nucleosome remodeling. eLife 5.

Brzezinka, K., Altmann, S., and Bäurle, I. (2019). BRUSHY1/TONSOKU/MGOUN3 is required for heat stress memory. Plant Cell Environ 42:771–781.

Burger, L., Gaidatzis, D., Schübeler, D., and Stadler, M. B. (2013). Identification of active regulatory regions from DNA methylation data. Nucleic Acids Res 41:e155.

Chang, C. Y., Lin, W. D., and Tu, S. L. (2014). Genome-wide analysis of heat-sensitive alternative splicing in Physcomitrella patens. Plant Physiology 165:826–840.

Charng, Y. Y., Liu, H. C., Liu, N. Y., Hsu, F. C., and Ko, S. S. (2006). Arabidopsis Hsa32, a novel heat shock protein, is essential for acquired thermotolerance during long recovery after acclimation. Plant Physiology 140:1297–1305.

Charng, Y.-Y., Liu, H.-C., Liu, N.-Y., Chi, W.-T., Wang, C.-N., Chang, S.-H., and Wang, T.-T. (2007). A heat-inducible transcription factor, HsfA2, is required for extension of acquired thermotolerance in Arabidopsis. Plant Physiol 143:251–262.

Correia, B., Valledor, L., Meijón, M., Rodriguez, J. L., Dias, M. C., Santos, C., Cañal, M. J., Rodriguez, R., and Pinto, G. (2013). Is the Interplay between Epigenetic Markers Related to the Acclimation of Cork Oak Plants to High Temperatures? PLOS ONE 8:e53543.

De Quattro, C., Mica, E., Pè, M. E., and Bertolini, E. (2018). Brachypodium distachyon Long Noncoding RNAs: Genome-Wide Identification and Expression Analysis. Methods Mol Biol 1667:31–42.

Ding, Y., Fromm, M., and Avramova, Z. (2012). Multiple exposures to drought “train” transcriptional responses in Arabidopsis. Nat Commun 3:740.

Ding, Y., Ma, Y., Liu, N., Xu, J., Hu, Q., Li, Y., Wu, Y., Xie, S., Zhu, L., Min, L., et al. (2017). microRNAs involved in auxin signalling modulate male sterility under high-temperature stress in cotton (Gossypium hirsutum). Plant J 91:977–994.

Feldeverd, E., Porter, B. W., Yuen, C. Y. L., Iwai, K., Carrillo, R., Smith, T., Barela, C., Wong, K., Wang, P., Kang, B.-H., et al. (2020). The Arabidopsis Protein Disulfide Isomerase Subfamily M Isoform, PDI9, Localizes to the Endoplasmic Reticulum and Influences Pollen Viability and Proper Formation of the Pollen Exine During Heat Stress. Frontiers in Plant Science 11.

Feng, H., and Wu, H. (2019). Differential methylation analysis for bisulfite sequencing using DSS. Quant Biol 7:327–334.

Feng, X. J., Li, J. R., Qi, S. L., Lin, Q. F., Jin, J. B., and Hua, X. J. (2016). Light affects salt stress-induced transcriptional memory of *P5CS1* in *Arabidopsis*. Proc. Natl. Acad. Sci. USA 113.

Ferrari, C., and Mutwil, M. (2019). Gene expression analysis of Cyanophora paradoxa reveals conserved abiotic stress responses between basal algae and flowering plants. New Phytologist Advance Access published 2019, doi:10.1111/nph.16257.

Friedrich, T., Oberkofler, V., Trindade, I., Altmann, S., Brzezinka, K., Lämke, J., Gorka, M., Kappel, C., Sokolowska, E., Skirycz, A., et al. (2021). Heteromeric HSFA2/HSFA3 complexes drive transcriptional memory after heat stress in Arabidopsis. Nat Commun 12:3426.

Garcia-Molina, A., Kleine, T., Schneider, K., Mühlhaus, T., Lehmann, M., and Leister, D. (2020). Translational Components Contribute to Acclimation Responses to High Light, Heat, and Cold in Arabidopsis. iScience 23:101331.

Giacomelli, J. I., Weigel, D., Chan, R. L., and Manavella, P. A. (2012). Role of recently evolved miRNA regulation of sunflower HaWRKY6 in response to temperature damage. New Phytol 195:766–773.

Goodstein, D. M., Shu, S., Howson, R., Neupane, R., Hayes, R. D., Fazo, J., Mitros, T., Dirks, W., Hellsten, U., Putnam, N., et al. (2012). Phytozome: A comparative platform for green plant genomics. Nucleic Acids Research 40.

Guan, Q., Lu, X., Zeng, H., Zhang, Y., and Zhu, J. (2013). Heat stress induction of miR398 triggers a regulatory loop that is critical for thermotolerance in Arabidopsis. Plant J 74:840–851.

Hansen, B. O., Vaid, N., Musialak-Lange, M., Janowski, M., and Mutwil, M. (2014). Elucidating gene function and function evolution through comparison of co-expression networks of plants. Frontiers in Plant Science 5.

He, X., Guo, S., Wang, Y., Wang, L., Shu, S., and Sun, J. (2020). Systematic identification and analysis of heat-stress-responsive lncRNAs, circRNAs and miRNAs with associated co-expression and ceRNA networks in cucumber (Cucumis sativus L.). Physiol Plant 168:736–754.

Hossain, M. S., Kawakatsu, T., Kim, K. D., Zhang, N., Nguyen, C. T., Khan, S. M., Batek, J. M., Joshi, T., Schmutz, J., Grimwood, J., et al. (2017). Divergent cytosine DNA methylation patterns in single-cell, soybean root hairs. New Phytol 214:808–819.

Hu, C., Lin, S., Chi, W., and Charng, Y. (2012). Recent gene duplication and subfunctionalization produced a mitochondrial GrpE, the nucleotide exchange factor of the Hsp70 complex, specialized in thermotolerance to chronic heat stress in Arabidopsis. Plant Physiol 158:747–758.

Hu, Z., Song, N., Zheng, M., Liu, X., Liu, Z., Xing, J., Ma, J., Guo, W., Yao, Y., Peng, H., et al. (2015). Histone acetyltransferase GCN5 is essential for heat stress-responsive gene activation and thermotolerance in Arabidopsis. Plant J 84:1178–1191.

Huang, Y., An, J., Sircar, S., Bergis, C., Lopes, C. D., He, X., Da Costa, B., Tan, F.-Q., Bazin, J., Antunez-Sanchez, J., et al. (2023). HSFA1a modulates plant heat stress responses and alters the 3D chromatin organization of enhancer-promoter interactions. Nat Commun 14:469.

Ito, H., Gaubert, H., Bucher, E., Mirouze, M., Vaillant, I., and Paszkowski, J. (2011). An siRNA pathway prevents transgenerational retrotransposition in plants subjected to stress. Nature 472:115–119.

Jaskiewicz, M., Conrath, U., and Peterhänsel, C. (2011). Chromatin modification acts as a memory for systemic acquired resistance in the plant stress response. EMBO Rep 12:50–55.

Jha, U. C., Nayyar, H., Jha, R., Khurshid, M., Zhou, M., Mantri, N., and Siddique, K. H. M. (2020). Long noncoding RNAs: emerging players regulating plant abiotic stress response and adaptation. BMC Plant Biology 20:466.

Kim, J.-M., To, T. K., Ishida, J., Matsui, A., Kimura, H., and Seki, M. (2012). Transition of chromatin status during the process of recovery from drought stress in Arabidopsis thaliana. Plant Cell Physiol 53:847–856.

Kim, D., Pertea, G., Trapnell, C., Pimentel, H., Kelley, R., and Salzberg, S. L. (2013). TopHat2: Accurate alignment of transcriptomes in the presence of insertions, deletions and gene fusions. Genome Biology 14.

Korotko, U., Chwiałkowska, K., Sa ko-Sawczenko, I., and Kwasniewski, M. (2021). DNA Demethylation in Response to Heat Stress in Arabidopsis thaliana. Int J Mol Sci 22:1555.

Kozomara, A., Birgaoanu, M., and Griffiths-Jones, S. (2019). miRBase: from microRNA sequences to function. Nucleic Acids Res 47:D155–D162.

Krueger, F., and Andrews, S. R. (2011). Bismark: a flexible aligner and methylation caller for Bisulfite-Seq applications. Bioinformatics 27:1571–1572.

Lämke, J., and Bäurle, I. (2017). Epigenetic and chromatin-based mechanisms in environmental stress adaptation and stress memory in plants.

Lämke, J., Brzezinka, K., Altmann, S., and Bäurle, I. (2016). A hit-and-run heat shock factor governs sustained histone methylation and transcriptional stress memory. EMBO J 35:162–175.

Langmead, B. (2010). Aligning short sequencing reads with Bowtie. Curr Protoc Bioinformatics Chapter 11:Unit 11.7.

Larkindale, J., Hall, J. D., Knight, M. R., and Vierling, E. (2005). Heat stress phenotypes of Arabidopsis mutants implicate multiple signaling pathways in the acquisition of thermotolerance. Plant Physiol 138:882–897.

Lee, J.-H., Yoon, H.-J., Terzaghi, W., Martinez, C., Dai, M., Li, J., Byun, M.-O., and Deng, X. W. (2010). DWA1 and DWA2, Two Arabidopsis DWD Protein Components of CUL4-Based E3 Ligases, Act Together as Negative Regulators in ABA Signal Transduction. The Plant Cell 22:1716–1732.

Li, H., Handsaker, B., Wysoker, A., Fennell, T., Ruan, J., Homer, N., Marth, G., Abecasis, G., and Durbin, R. (2009). The Sequence Alignment/Map format and SAMtools. Bioinformatics 25:2078–2079.

Li, S., Liu, J., Liu, Z., Li, X., Wu, F., and He, Y. (2014). HEAT-INDUCED TAS1 TARGET1 Mediates Thermotolerance via HEAT STRESS TRANSCRIPTION FACTOR A1a-Directed Pathways in Arabidopsis. Plant Cell 26:1764–1780.

Li, J., Ma, W., Zeng, P., Wang, J., Geng, B., Yang, J., and Cui, Q. (2015). LncTar: a tool for predicting the RNA targets of long noncoding RNAs. Brief Bioinform 16:806–812.

Li, S., Castillo-González, C., Yu, B., and Zhang, X. (2017a). The functions of plant small RNAs in development and in stress responses. The Plant Journal 90:654–670.

Li, J., Labbadia, J., and Morimoto, R. I. (2017b). Rethinking HSF1 in Stress, Development, and Organismal Health. Trends Cell Biol 27:895–905.

Liao, Y., Smyth, G. K., and Shi, W. (2014). featureCounts: an efficient general purpose program for assigning sequence reads to genomic features. Bioinformatics 30:923–930.

Ling, Y., Serrano, N., Gao, G., Atia, M., Mokhtar, M., Woo, Y. H., Bazin, J., Veluchamy, A., Benhamed, M., Crespi, M., et al. (2018). Thermopriming triggers splicing memory in Arabidopsis. J Exp Bot 69:2659–2675.

Liu, J., and He, Z. (2020). Small DNA Methylation, Big Player in Plant Abiotic Stress Responses and Memory. Frontiers in Plant Science 11.

Liu, J., Feng, L., Li, J., and He, Z. (2015). Genetic and epigenetic control of plant heat responses. Frontiers in Plant Science 6.

Liu, T., Li, Y., Duan, W., Huang, F., and Hou, X. (2017). Cold acclimation alters DNA methylation patterns and confers tolerance to heat and increases growth rate in Brassica rapa. J Exp Bot 68:1213–1224.

Liu, H., Lämke, J., Lin, S., Hung, M.-J., Liu, K.-M., Charng, Y., and Bäurle, I. (2018). Distinct heat shock factors and chromatin modifications mediate the organ-autonomous transcriptional memory of heat stress. The Plant Journal 95:401–413.

Lobell, D. B., Schlenker, W., and Costa-Roberts, J. (2011). Climate trends and global crop production since 1980. Science 333:616–620.

Lohse, M., Nagel, A., Herter, T., May, P., Schroda, M., Zrenner, R., Tohge, T., Fernie, A. R., Stitt, M., and Usadel, B. (2014). Mercator: A fast and simple web server for genome scale functional annotation of plant sequence data. Plant, Cell and Environment 37:1250–1258.

López, M.-E., Roquis, D., Becker, C., Denoyes, B., and Bucher, E. (2022). DNA methylation dynamics during stress response in woodland strawberry (Fragaria vesca). Hortic Res 9:uhac174.

Love, M. I., Huber, W., and Anders, S. (2014). Moderated estimation of fold change and dispersion for RNA-seq data with DESeq2. Genome Biology 15.

Meng, X., Li, A., Yu, B., and Li, S. (2021). Interplay between miRNAs and lncRNAs: Mode of action and biological roles in plant development and stress adaptation. Computational and Structural Biotechnology Journal 19:2567–2574.

Min, L., Li, Y., Hu, Q., Zhu, L., Gao, W., Wu, Y., Ding, Y., Liu, S., Yang, X., and Zhang, X. (2014). Sugar and auxin signaling pathways respond to high-temperature stress during anther development as revealed by transcript profiling analysis in cotton. Plant Physiol 164:1293–1308.

Mozgová, I., Wildhaber, T., Liu, Q., Abou-Mansour, E., L’Haridon, F., Métraux, J.-P., Gruissem, W., Hofius, D., and Hennig, L. (2015). Chromatin assembly factor CAF-1 represses priming of plant defence response genes. Nature Plants 1:1–8.

Oberkofler, V., Pratx, L., and Bäurle, I. (2021). Epigenetic regulation of abiotic stress memory: maintaining the good things while they last. Current Opinion in Plant Biology 61:102007.

Ohama, N., Sato, H., Shinozaki, K., and Yamaguchi-Shinozaki, K. (2017). Transcriptional Regulatory Network of Plant Heat Stress Response. Trends in Plant Science 22:53–65.

Olas, J. J., Apelt, F., Annunziata, M. G., John, S., Richard, S. I., Gupta, S., Kragler, F., Balazadeh, S., and Mueller-Roeber, B. (2021). Primary carbohydrate metabolism genes participate in heat-stress memory at the shoot apical meristem of Arabidopsis thaliana. Mol Plant 14:1508–1524.

Pertea, G., and Pertea, M. (2020). GFF Utilities: GffRead and GffCompare. F1000Res 9:ISCB Comm J-304.

Pertea, M., Pertea, G. M., Antonescu, C. M., Chang, T.-C., Mendell, J. T., and Salzberg, S. L. (2015). StringTie enables improved reconstruction of a transcriptome from RNA-seq reads. Nat Biotechnol 33:290–295.

Popova, O. V., Dinh, H. Q., Aufsatz, W., and Jonak, C. (2013). The RdDM pathway is required for basal heat tolerance in Arabidopsis. Mol Plant 6:396–410.

Richter, K., Haslbeck, M., and Buchner, J. (2010). The heat shock response: life on the verge of death. Mol Cell 40:253–266.

Scafaro, A. P., Fan, Y., Posch, B. C., Garcia, A., Coast, O., and Atkin, O. K. (2021). Responses of leaf respiration to heatwaves. Plant Cell Environ 44:2090–2101.

Sedaghatmehr, M., Mueller-Roeber, B., and Balazadeh, S. (2016). The plastid metalloprotease FtsH6 and small heat shock protein HSP21 jointly regulate thermomemory in Arabidopsis. Nat Commun 7:12439.

Sedaghatmehr, M., Stüwe, B., Mueller-Roeber, B., and Balazadeh, S. (2022). Heat shock factor HSFA2 fine-tunes resetting of thermomemory via plastidic metalloprotease FtsH6. J Exp Bot 73:6394–6404.

Shen, S., Park, J. W., Lu, Z. X., Lin, L., Henry, M. D., Wu, Y. N., Zhou, Q., and Xing, Y. (2014a). rMATS: Robust and flexible detection of differential alternative splicing from replicate RNA-Seq data. Proceedings of the National Academy of Sciences of the United States of America 111:E5593–E5601.

Shen, X., Jonge, J. D., Forsberg, S. K. G., Pettersson, M. E., Sheng, Z., Hennig, L., and Carlborg, Ö. (2014b). Natural CMT2 Variation Is Associated With Genome-Wide Methylation Changes and Temperature Seasonality. PLOS Genetics 10:e1004842.

Shiva, S., Samarakoon, T., Lowe, K. A., Roach, C., Vu, H. S., Colter, M., Porras, H., Hwang, C., Roth, M. R., Tamura, P., et al. (2020). Leaf Lipid Alterations in Response to Heat Stress of Arabidopsis thaliana. Plants (Basel) 9:845.

Singh, P., Yekondi, S., Chen, P.-W., Tsai, C.-H., Yu, C.-W., Wu, K., and Zimmerli, L. (2014). Environmental History Modulates Arabidopsis Pattern-Triggered Immunity in a HISTONE ACETYLTRANSFERASE1-Dependent Manner. Plant Cell 26:2676–2688.

Song, Y., Ci, D., Tian, M., and Zhang, D. (2016). Stable methylation of a noncoding RNA gene regulates gene expression in response to abiotic stress in Populus simonii. J Exp Bot 67:1477–1492.

Song, X., Li, Y., Cao, X., and Qi, Y. (2019). MicroRNAs and Their Regulatory Roles in Plant-Environment Interactions. Annu Rev Plant Biol 70:489–525.

Song, X., Hu, J., Wu, T., Yang, Q., Feng, X., Lin, H., Feng, S., Cui, C., Yu, Y., Zhou, R., et al. (2021). Comparative analysis of long noncoding RNAs in angiosperms and characterization of long noncoding RNAs in response to heat stress in Chinese cabbage. Hortic Res 8:1–21.

Stief, A., Altmann, S., Hoffmann, K., Pant, B. D., Scheible, W. R., and Bäurle, I. (2014). Arabidopsis miR156 regulates tolerance to recurring environmental stress through SPL transcription factors. Plant Cell 26:1792–1807.

Tan, Q. W., Lim, P. K., Chen, Z., Pasha, A., Provart, N., Arend, M., Nikoloski, Z., and Mutwil, M. (2023). Cross-stress gene expression atlas of Marchantia polymorpha reveals the hierarchy and regulatory principles of abiotic stress responses. Nat Commun 14:986.

Wang, A., Hu, J., Gao, C., Chen, G., Wang, B., Lin, C., Song, L., Ding, Y., and Zhou, G. (2019). Genome-wide analysis of long noncoding RNAs unveils the regulatory roles in the heat tolerance of Chinese cabbage (Brassica rapa ssp.chinensis). Sci Rep 9:5002.

Xin, M., Wang, Y., Yao, Y., Song, N., Hu, Z., Qin, D., Xie, C., Peng, H., Ni, Z., and Sun, Q. (2011). Identification and characterization of wheat long non-protein coding RNAs responsive to powdery mildew infection and heat stress by using microarray analysis and SBS sequencing. BMC Plant Biology 11:61.

Yadav, V. K., Singh, S., Yadav, A., Agarwal, N., Singh, B., Jalmi, S. K., Yadav, V. K., Tiwari, V. K., Kumar, V., Singh, R., et al. (2022). Stress Conditions Modulate the Chromatin Interactions Network in Arabidopsis. Frontiers in Genetics 12.

Yang, L., Wu, L., Chang, W., Li, Z., Miao, M., Li, Y., Yang, J., Liu, Z., and Tan, J. (2018). Overexpression of the maize E3 ubiquitin ligase gene ZmAIRP4 enhances drought stress tolerance in Arabidopsis. Plant Physiol Biochem 123:34–42.

Yu, X., Yang, J., Li, X., Liu, X., Sun, C., Wu, F., and He, Y. (2013). Global analysis of cis-natural antisense transcripts and their heat-responsive nat-siRNAs in Brassica rapa. BMC Plant Biol 13:208.

Zhao, J., He, Q., Chen, G., Wang, L., and Jin, B. (2016). Regulation of noncoding RNAs in heat stress responses of plants. Frontiers in Plant Science 7.

Zhao, J., Lu, Z., Wang, L., and Jin, B. (2020). Plant Responses to Heat Stress: Physiology, Transcription, Noncoding RNAs, and Epigenetics. Int J Mol Sci 22:117.

Zhou, J., Zhang, Y., Qi, J., Chi, Y., Fan, B., Yu, J.-Q., and Chen, Z. (2014). E3 Ubiquitin Ligase CHIP and NBR1-Mediated Selective Autophagy Protect Additively against Proteotoxicity in Plant Stress Responses. PLOS Genetics 10:e1004116.

Zilberman, D., Gehring, M., Tran, R. K., Ballinger, T., and Henikoff, S. (2007). Genome-wide analysis of Arabidopsis thaliana DNA methylation uncovers an interdependence between methylation and transcription. Nat Genet 39:61–69.

