## Supplementary material for "Multi-omic dissection of ancestral heat stress memory responses in *Brachypodium distachyon*": Figure S1-6

### Figure S1. Proof for experimentally establishing the occurrence of heat stress memory in *Brachypodium distachyon*.

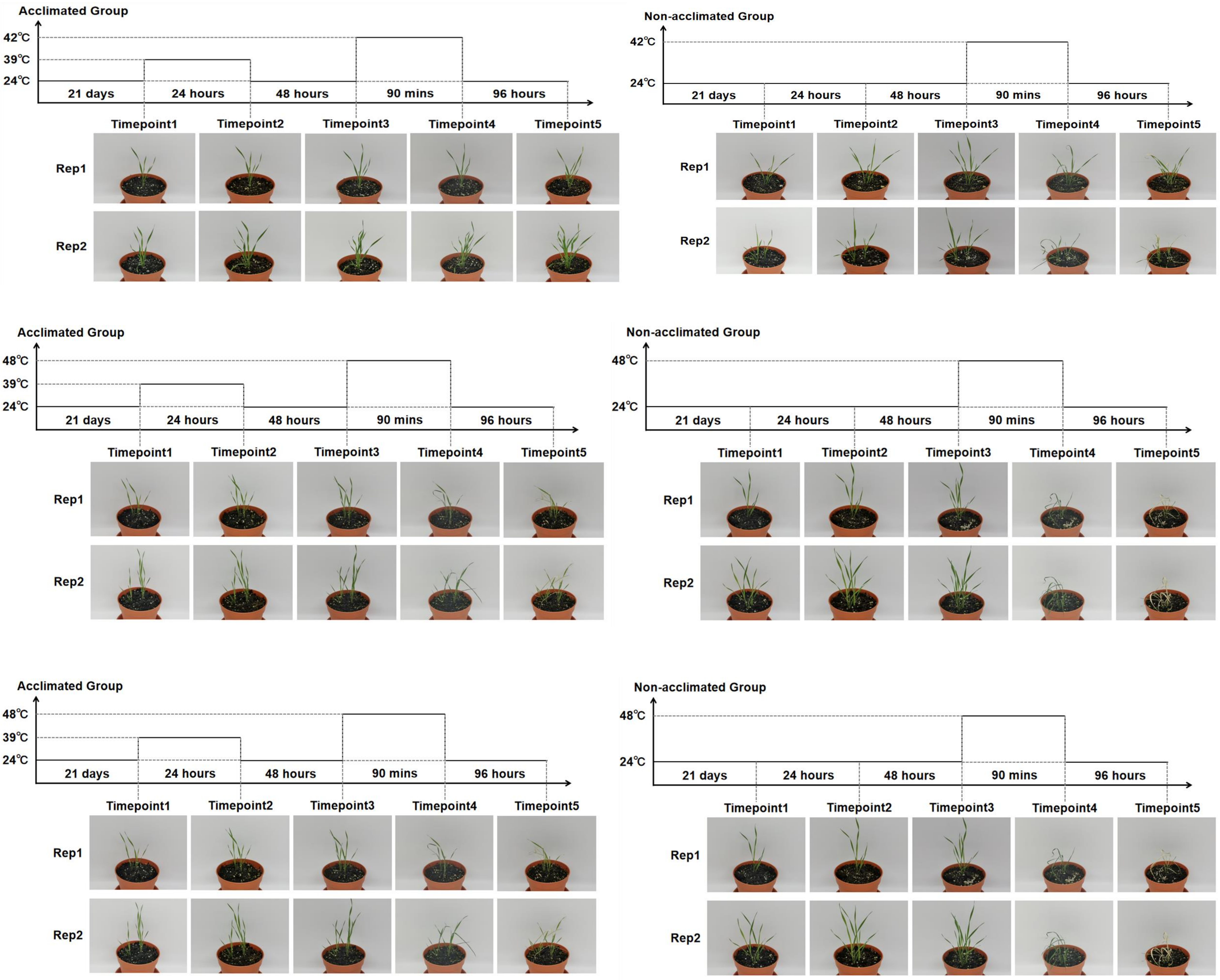

**Figure S2. Pipeline for filtering lncRNAs from assembled transcripts.**

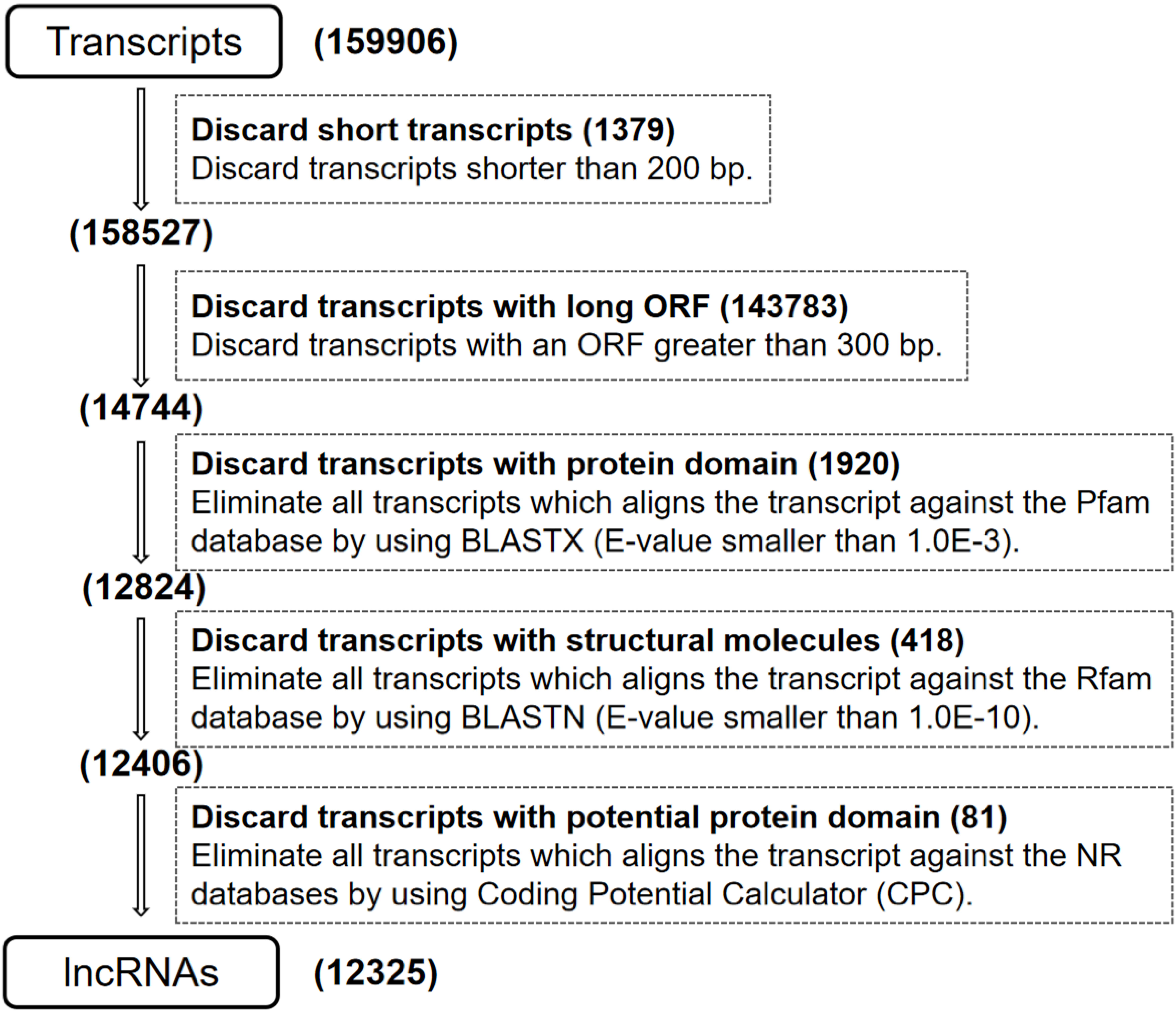

**Figure S3. Length distribution of identified lncRNAs.**

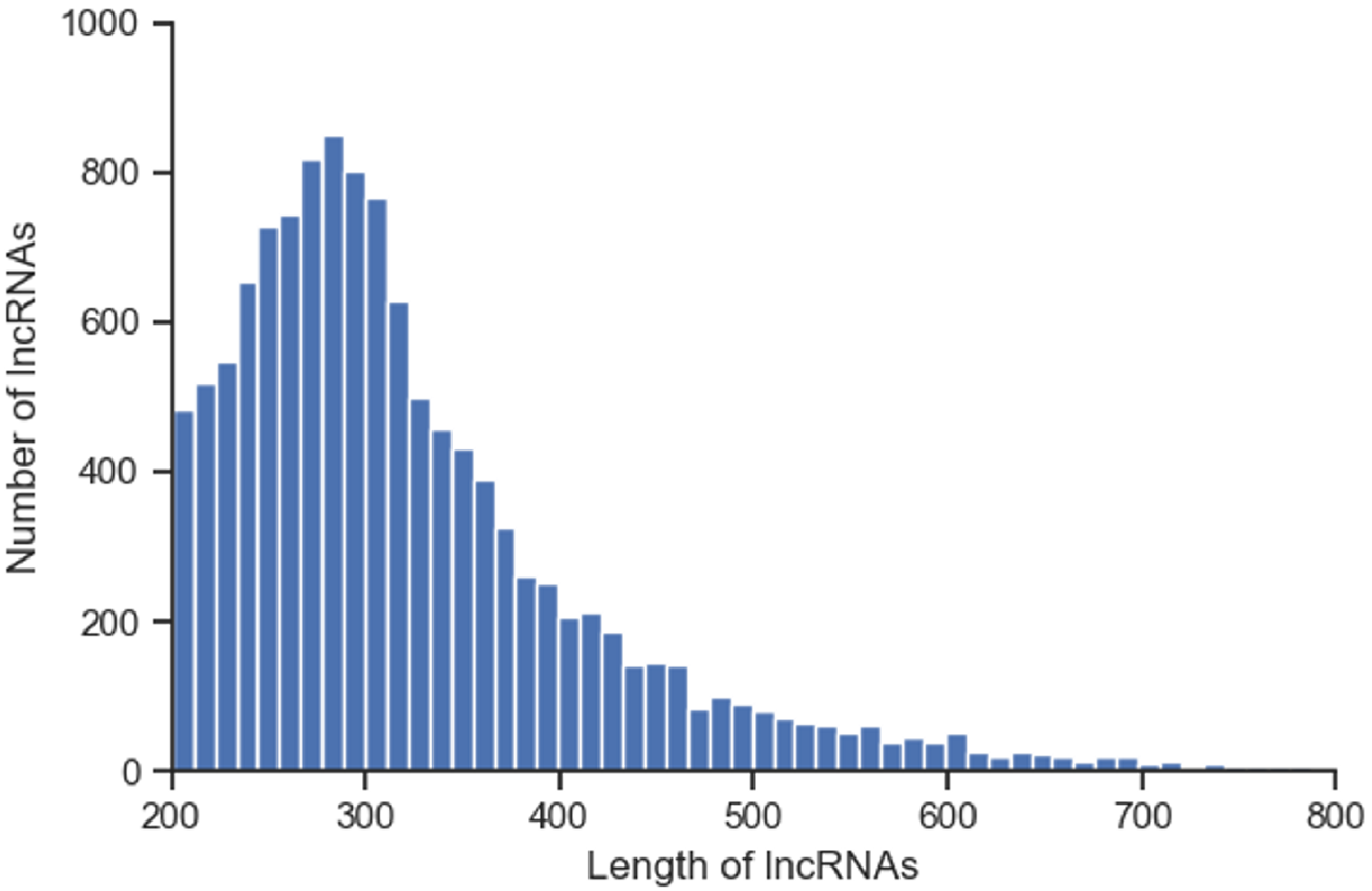

### Figure S4

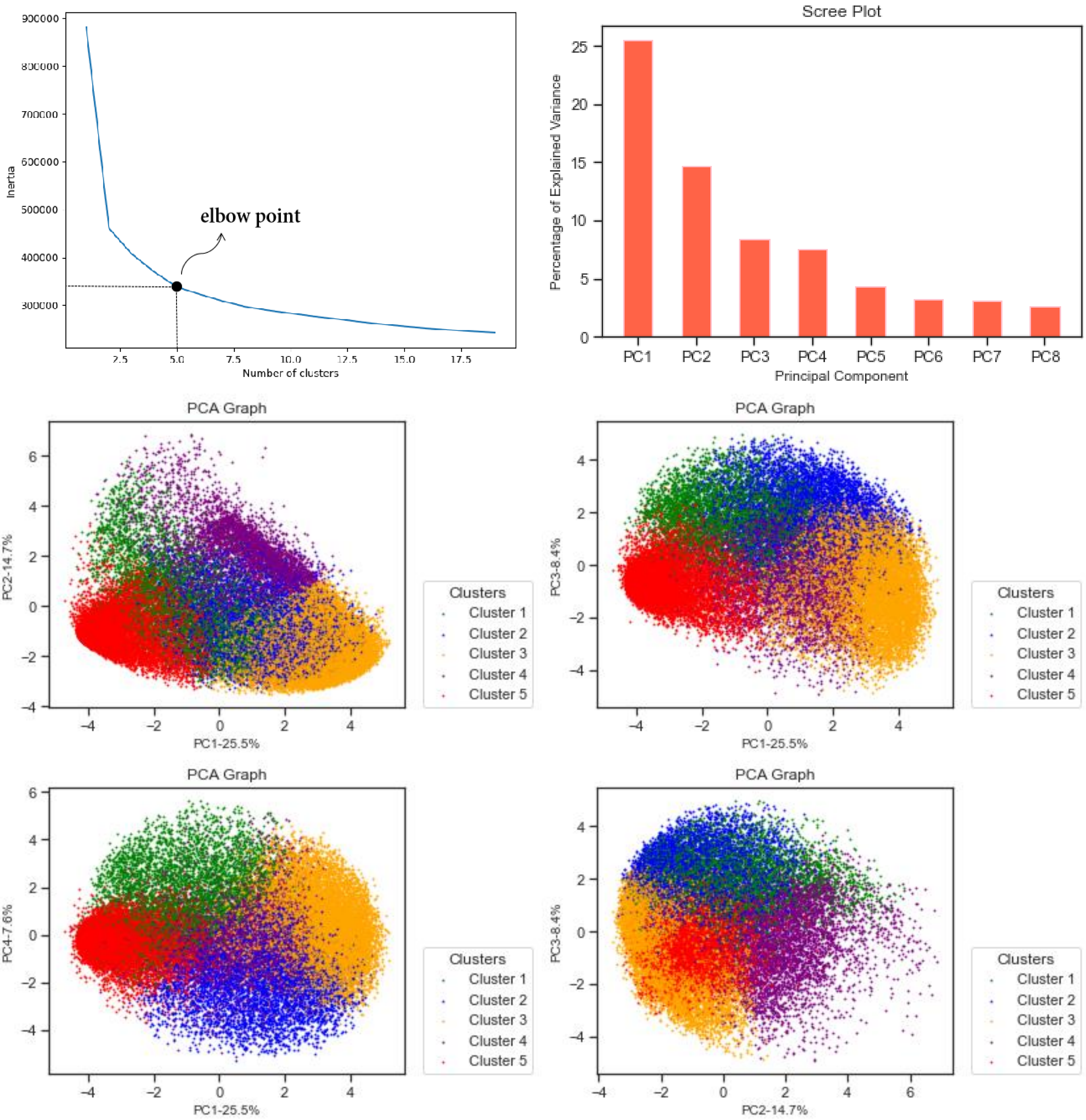

elbow point
Inertia
Number of clusters
Scree Plot
Percentage of Explained Variance
Principal Component
PC1
PC2
PC3
PC4
PC5
PC6
PC7
PC8
PCA Graph
PC2-14.7%
PC1-25.5%
PC3-8.4%
PC4-7.6%
Clusters
Cluster 1
Cluster 2
Cluster 3
Cluster 4
Cluster 5

### Figure S5. UpsetR plot for ASGs and DASGs from AllvsAA1 comparisons.

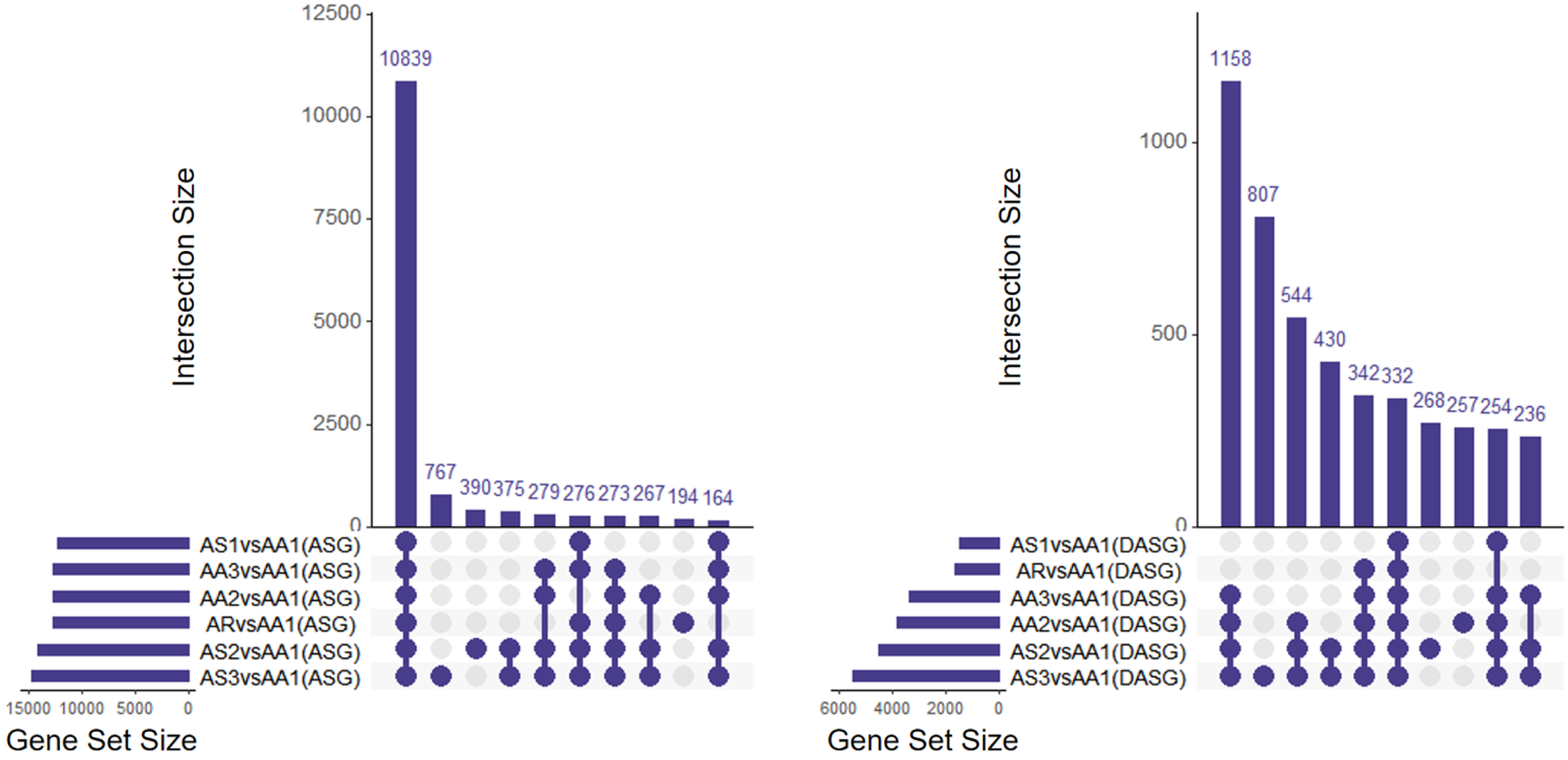


**ASG** **DAST**

### Figure S6. Distribution of segments on a 2-D plot displaying the level of methylation and the number of CpGs.

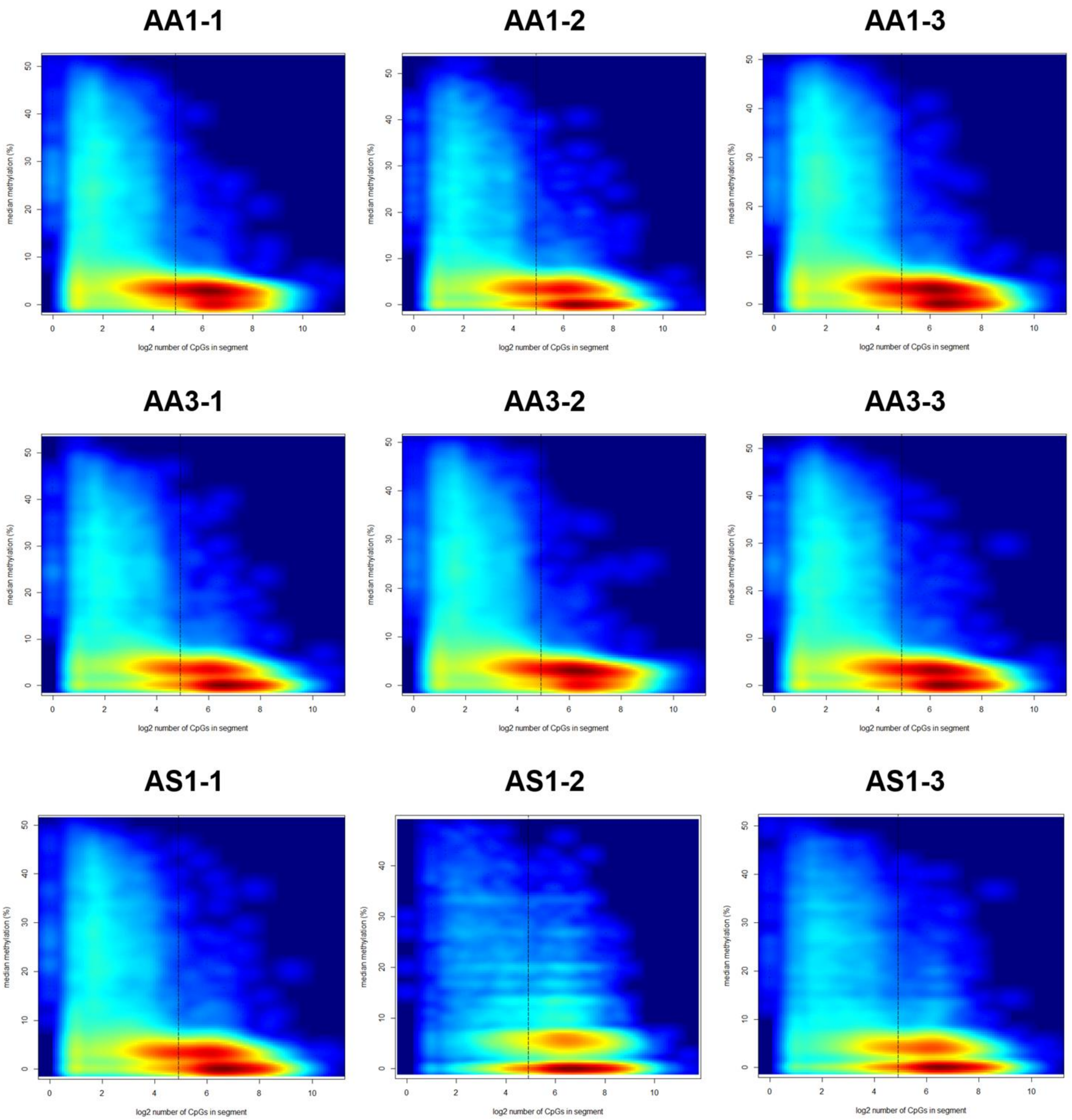
